# Fast A-type currents shape a rapidly adapting form of delayed short latency firing of excitatory superficial dorsal horn neurons that express the NPY Y1 receptor

**DOI:** 10.1101/2021.02.18.431823

**Authors:** Ghanshyam P. Sinha, Pranav Prasoon, Bret N. Smith, Bradley K. Taylor

## Abstract

Neuroanatomical and behavioral evidence indicates that neuropeptide Y Y1 receptor-expressing interneurons (Y1-INs) in the superficial dorsal horn (SDH) are predominantly excitatory and contribute to chronic pain. Using an adult *ex vivo* spinal cord slice preparation from Y1eGFP reporter mice, we characterized firing patterns in response to steady state depolarizing current injection of GFP-positive cells in lamina II, the great majority of which expressed Y1 mRNA (88%). Randomly sampled and Y1eGFP neurons exhibited five firing patterns: tonic (TF), initial burst (IBF), phasic (PF), delayed short-latency <180 ms (DSLF), and delayed long-latency >180 ms (DLLF). When studied at resting membrane potential, most RS neurons exhibited delayed firing, while most Y1eGFP neurons exhibited phasic firing and not delayed firing. A preconditioning membrane hyperpolarization produced only subtle changes in the firing patterns of randomly sampled neurons, but dramatically shifted Y1eGFP neurons to DSLF (46%) and DLLF (24%). In contrast to randomly sampled DSLF neurons which rarely exhibited spike frequency adaptation, Y1eGFP DSLF neurons were almost always rapidly adapting, a characteristic of nociceptive-responsive SDH neurons. Rebound spiking was more prevalent in Y1eGFP neurons (6% RS vs 32% Y1eGFP), indicating enrichment of T-type calcium currents. Y1eGFP DSLF neurons exhibited fast A-type potassium currents that are known to delay or limit action potential firing, and these were of smaller current density as compared to randomly sampled DSLF neurons. Our results inspire future studies to determine whether tissue or nerve injury downregulates channels that contribute to A-currents, thus potentially unmasking T-type calcium channel activity and membrane hyperexcitability in Y1-INs, leading to persistent pain.

**KEYPOINTS:**

- Neuropeptide Y Y1 receptor-expressing neurons in the dorsal horn of the spinal cord contribute to chronic pain.
- For the first time, we characterized the firing patterns of Y1-expressing neurons in Y1eGFP reporter mice.
- Under hyperpolarized conditions, most Y1eGFP neurons exhibited fast A-type potassium currents and delayed, short-latency firing (DSLF).
- Y1eGFP DSLF neurons were almost always rapidly adapting and often exhibited rebound spiking, characteristics of spinal pain neurons under the control of T-type calcium channels.
- These results inspire future studies to determine whether tissue or nerve injury downregulates the channels that underlie A-currents, thus unmasking membrane hyperexcitability in Y1- expressing dorsal horn neurons, leading to persistent pain

## INTRODUCTION

Neuropeptide Y (NPY) Y1 receptor-expressing interneurons (Y1-INs) are widely distributed in the spinal cord, with the greatest density in lamina II of the dorsal horn (Brumovsky *et al*., 2006; Shi *et al*., 2006), a region that is critical for the spinal modulation of noxious somatosensory input (Todd & Koerber, 2006; Todd, 2017). Neuroanatomical and behavioral evidence indicates that these Y1-INs are predominantly excitatory and contribute to chronic neuropathic pain (Naveilhan *et al*., 2001; Smith *et al*., 2007; Intondi *et al*., 2008; Wiley *et al*., 2009; Solway *et al*., 2011: Nelson et al., 2019). Despite this, the electrophysiological characteristics of Y1-INs remain largely undefined beyond a small number of studies that identified putative Y1-INs based on their responsiveness to NPY agonists (Moran *et al*., 2004; Miyakawa *et al*., 2005; Melnick, 2012). Whole cell recordings in the rat spinal cord slice consistently indicate that bath application of NPY inhibits both presynaptic and postsynaptic components of excitatory neurotransmission in the dorsal horn, and the prevailing view is that Y1 mediates the ability of NPY to post-synaptically inhibit spinal interneurons. For example, bath administration of NPY or the Y1 receptor-specific agonist [Leu^31^,Pro^34^]-NPY consistently produced an outward current and membrane hyperpolarization in spinal interneurons that could be blocked with Y1 but not Y2 antagonists (Miyakawa *et al*., 2005; Melnick, 2012). Furthermore, NPY or [Leu31,Pro34]-NPY decreased noxious stimulus-and dorsal root stimulus-evoked substance P release into the dorsal horn (Taylor *et al*., 2014). However, further studies are needed to profile subsets of Y1-INs based on molecular, morphological, and/or electrophysiological characteristics. One exciting approach uses transgenic reporter mice to better elucidate intrinsic and evoked firing patterns and perhaps reveal functionally distinct subclasses of Y1-INs (Smith *et al*., 2015; Dickie *et al*., 2019).

Rigorous classification of dorsal horn neurons is an essential step towards a better understanding of the complex microcircuitry that determines spinal pain transmission. Classification systems are based on morphology, neurochemical identity, and neuronal firing patterns (Todd & Spike, 1993; Todd & Koerber, 2006). For example, recordings of action potential firing patterns in response to current injection has provided comprehensive classification schemes to identify neurons with similar input-output functions (Prescott & De Koninck, 2002; Ruscheweyh & Sandkühler, 2002; Ruscheweyh *et al*., 2004; Heinke *et al*., 2004; Schoffnegger *et al*., 2006; Yasaka *et al*., 2010; Li & Baccei, 2014; Punnakkal *et al*., 2014; Smith *et al*., 2015; Stebbing *et al*., 2016). These properties have yielded greater understanding of the mechanisms that underlie the processing and integration of spinal nociceptive signals (Prescott & De Koninck, 2002; Grudt & Perl, 2002; Ruscheweyh & Sandkühler, 2002; Balachandar & Prescott, 2018). Voltage-clamp recordings provide additional information about ion channels underlying specific neural activity and have been used to further refine this firing pattern-based classification of dorsal horn neurons (Ruscheweyh & Sandkühler, 2002; Ruscheweyh *et al*., 2004; Yasaka *et al*., 2010; Smith *et al*., 2015), leading to a better understanding of neuronal plasticity after injury, and guiding the development of novel therapeutic targets for chronic pain (Ashcroft, 2000). Examples include electrophysiological studies promoting Cav3.2 channels (Candelas *et al*., 2019) and channels mediating I_h_ currents (Tsantoulas *et al*., 2017) as targets for pain therapeutics.

Here, we tested the hypothesis that neuropeptide Y (NPY) Y1 receptor expressing (Y1R) neurons exhibit firing properties and underlying voltage-gated currents that distinguish them from the overall SDH population. We assessed whether the Y1R neurons are a subpopulation of the heterogeneous spinal superficial dorsal horn (SDH) neurons that express a wide variety of morphologies, firing patterns, and neurochemical markers (Todd, 2010). We first confirmed that GFP-labelled neurons from Y1eGFP reporter mice expressed Y1 mRNA and can respond to NPY. We next compared their firing patterns to randomly sampled SDH neurons, and then identified voltage activated currents that underlie specific activity patterns.

## METHODS

### Ethical approval

All mice were cared for in accordance with the Guide for the Care and Use of Laboratory Animals. Animals were group-housed in a well-ventilated facility under standard housing conditions (12:12 h light: dark cycle with lights on at 7:00 a.m. and temperatures of 20° to 22°C with *ad libitum* access to water and food) and breeding colonies were maintained in-house. Male and female mice aged 4-10 weeks were used. There were no differences in results that were disaggregated by sex, so results from both sexes are pooled. All animal use protocols were approved by the Institutional Animal Care and Use Committee of the University of Kentucky (protocol number 2017-2731) and/or the University of Pittsburgh (protocol number 18062960).

### Transgenic mice

BAC transgenic mice expressing enhanced green fluorescent protein (eGFP) under the control of the Y1R promoter (i.e., Y1ReGFP mice, RRID:MMRRC_010554-UCD) used in this study were generated by the Gene Expression Nervous System Atlas (GENSAT) project (Heintz, 2004). They were backcrossed to a Swiss Webster background and maintained by heterozygote x homozygote breeding, yielding transgenic (+/-) mice for Y1ReGFP neuronal recordings and wildtype (+/+) control littermates for randomly sampled neuronal recordings.

### Combined Immunohistochemistry and Fluorescence in situ Hybridization (FISH)

Currently available antibodies directed at the Y1 receptor do not yield specific staining in the mouse spinal cord. As an alternative, we used *in situ* hybridization to localize Y1-mRNA (RNAscope Multiplex Fluorescent V2 Assay, Advanced Cell Diagnostics, Hayward, CA, USA) in sections from Y1eGFP mice. Y1eGFP mice at 6-8 weeks of age were transcardially perfused with chilled phosphate-buffered saline (PBS) followed by 4% paraformaldehyde (Sigma-Aldrich, USA) in PBS. The lumbar spinal cord (L3-L5) was removed, post-fixed in 4% paraformaldehyde for 4 hr, cryoprotected overnight in 15% sucrose followed by 30% sucrose in PBS, and then embedded (OCT, Tissue Tek). Tissues were sectioned on a cryostat at 8 µm and mounted onto slides (Superfrost Plus, Fisher Scientific). Slides were rinsed with PBS, pretreated for 15 min with protease per manufacturer’s instructions, and then incubated and hybridized with Y1-mRNA probe for 2 hr at 40° C in a HybEZ oven (Advanced Cell Diagnostics, Hayward, CA, USA). Amplification and detection steps were performed using the RNAscope Multiplex Fluorescent Reagent Kit v2 (Advanced Cell Diagnostics, Hayward CA, USA; Cat. No. 323100), three proprietary signal amplification system (Amp1, Amp2, and Amp3 to detect multiple target RNA signals) and proprietary wash buffers. Sections were incubated with Amp1 for 30 min at 40°C and then washed three times with wash buffer for 2 min each. The next incubation was with Amp 2 for 30 min at 40 °C, followed by three washes with wash buffer for 2 min each. Lastly, sections were incubated with Amp 3 for 15 min at 40 °C and washed three times with wash buffer for 2 min each. In the last step, sections were incubated with appropriate HRP channel and TSA Plus fluorophore reagent for 15 and 30 min respectively at 40 °C and washed three times with wash buffer for 2 min each. Because the tissue heating steps required for hybridization diminished the GFP signal, sections were incubated with anti-GFP primary antibody (Chicken,1:1000; Abcam) overnight at 4° C on a slow rocker, followed by 3 washes with PBS and incubation with secondary antibody (Alexa Fluor 488 goat anti-chicken, 1:1000; Invitrogen) on a slow rocker at room temperature for 60 min, washed with PBS followed by another wash with 0.01-M phosphate buffer, then air-dried and cover-slipped with Hard Set Antifade Mounting Media with DAPI (Vectashield; Vector Labs). Sections were then imaged on a Nikon Eclipse Ti2 microscope; images captured at 20x were analyzed using NIS-Elements Advanced Research software v5.02.

### Quantification of co-localization of Y1eGFP and Y1-mRNA profiles

We focused our quantification of co-localization within lamina I-III, where most nociceptive peripheral afferents terminate within the dorsal horn (Todd, 2010). For quantification of in situ hybridization, only soma containing 3 or more puncta were counted as positive for Y1-mRNA. Y1eGFP-positive and Y1-mRNA-positive profiles were considered for quantification only if they contained DAPI-labeled nuclei. The number of Y1eGFP labelled and Y1-mRNA profiles were manually counted for each section averaged over 6-7 sections per animal.

### Slice Preparation for Electrophysiology

Mice were anesthetized with 5% isoflurane and perfused transcardially with 10 ml of ice-cold, sucrose-containing artificial cerebrospinal fluid (aCSF) (sucrose-aCSF) that contained (in mM): KCl 2.5, KH_2_PO_4_ 1.0, CaCl_2_ 1, MgSO_4_ 2.5, NaHCO_3_ 26, glucose 11, sucrose 235, oxygenated with 95% O_2_, 5% CO_2_. The lumbar spinal cord was rapidly isolated by laminectomy, placed in oxygenated, ice-cold, sucrose-aCSF, cleaned of dura mater, and nerve roots were cut close to the cord. The spinal cord was immersed in low-melting-point agarose (3% in sucrose-aCSF; Life Technologies, Carlsbad, CA, USA) and parasagittal slices (300–450 µm) were cut in ice-cold, sucrose-aCSF from one lateral side to the other side using a vibrating microtome (7000smz-2; Campden Instruments, Lafayette, IN, USA). All slices were incubated for 15 min at room temperature in recovery solution that contained (in mM): n-methyl-d-glucamine (NMDG) 92, KCl 2.5, NaH_2_PO_4_ 1.2, NaHCO_3_ 30, HEPES 20, glucose 25, sodium ascorbate 5, thiourea 2, sodium pyruvate 3, MgSO_4_ 10, CaCl_2_ 0.5 pH 7.3 (HCl). The slices were then transferred to normal aCSF used for recording, which contained (in mM): NaCl 126, KCl 2.5, NaH_2_PO_4_ 1.25, CaCl_2_ 2.0, MgSO_4_ 1.0, NaHCO_3_ 26, glucose 11, for an hour before beginning any experiment.

### Patch-clamp recordings

A single parasagittal spinal cord slice was transferred to a fixed stage mounted under an upright microscope (Olympus BX51 WI), where it was continuously superfused with oxygenated aCSF (Experimental setup: **Figure 1A**). Recordings from neurons were obtained under direct visualization using differential interference contrast (DIC) optics or with epifluorescent microscopy for Y1eGFP-positive cell bodies (Figure 1B). Recording pipettes (3-6 MΩ) containing (in mM): K-gluconate 135, NaCl 1, MgCl_2_ 2, CaCl_2_ 1, HEPES 5, EGTA 5, Mg-ATP 2 and Na_4_-GTP 0.2, pH 7.3 (∼300 mOsm/L). Patch-clamp recordings in current- and voltage-clamp modes were performed on SDH neurons using an Axon Instruments Multiclamp 700B amplifier (Molecular Devices). Signals were low-pass filtered at 4-20 kHz, amplified 1-20 fold, sampled at 5-10 kHz and analyzed offline using pClamp 10.3. Series resistance was less than 25 MΩ (mean=19±2 MΩ). No correction was made for the liquid junction potential (calculated value: -13.9 mV). Experimental data were recorded approximately 5-10 min after establishing whole-cell configuration. All recordings were performed at room temperature on neurons selected from the distinctive translucent band in the medial to mediolateral subdivisions of the SDH region, with a depth of roughly 20 to 100 µm (Yasaka *et al*., 2010).

**Figure 1.**
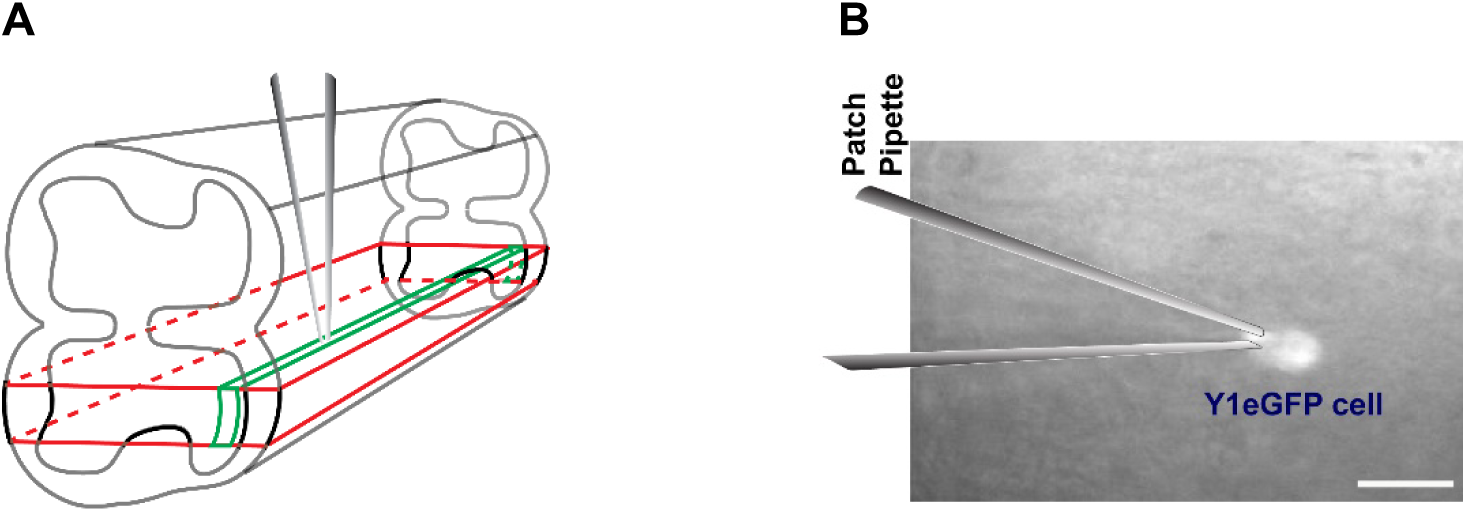
Experimental setup for electrophysiological recording. (**A**) Illustration of the parasagittal plane from which 300-450 µm slices were obtained from the lumbar L3/L4 segment of adult mouse spinal cord. (**B**) Fluorescent image of a recording pipette attached to a Y1eGFP cell within a para-sagittal slice. Scale bar – 10 µm.

### Firing patterns during current clamp recordings

Firing patterns were determined in response to a series of 7-9 depolarizing current injections (1 s duration) that were delivered every 8 s in increasing steps of 20 pA. Firing patterns were routinely elicited from three different conditioning membrane potentials: (a) between −50 and −65 mV; (b) between −65 and −80 mV (usually elicited from resting membrane potential (RMP)) and (c) between −80 and -95 mV to detect voltage-dependence of the firing patterns.

We calculated number of action potentials, latency to first action potential, initial spike frequency, average spike frequency, and spike frequency adaptation at Low, Medium, and High strengths of current injection (LCI, MCI and HCI, respectively) with algorithms written in MATLAB. LCI is the first current injection step that elicited AP firing. HCI represents the magnitude of current injection generating the maximum number of APs. MCI represents the magnitude of current injection that was in between (the average of) LCI and HCI. Firing frequency (f) was calculated as the reciprocal of the interspike interval (ISI) (Prescott & De Koninck, 2002). Firing adaptation was calculated as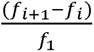, where *i* is the spike ordinate and f_1_ is the initial frequency (reciprocal of ISI between first and second spike). The latency to first action potential was measured from the onset of current injection to the time of first spike peak.

### Active membrane properties

The inflection point during spike initiation was used to determine the width and amplitude of the first action potential evoked by depolarizing current injected at resting membrane potential (Ruscheweyh *et al*., 2004; Schoffnegger *et al*., 2006). The amplitude of action potential afterhyperpolarization (AHP) was calculated as the difference from the AP inflection point to the maximum negative peak following the action potential. Action potential threshold (rheobase) was measured by means of a voltage step protocol that was applied to neurons held at −60 mV, with 5 ms voltage steps applied in 2 mV increments (from -50 mV towards depolarization) until the appearance of a fast Na^+^ current. We chose this method because it takes into account the facilitating or inhibiting effects of other voltage-dependent currents (like the A-type current) on the action potential generation (Ruscheweyh & Sandkühler, 2002; Ruscheweyh *et al*., 2004).

### Dorsal root stimulation (DRS)

Slices were prepared as described above, together with an approximately 4 mm long dorsal root entering the dorsal horn L3-L4 segments. Electrical stimulation of the attached dorsal root was performed using a glass suction electrode to evoke EPSCs (V_h_= -60). We used stimuli of 0.1 ms pulse width at 0.03 Hz and variable intensities (10 to 600 µA), which putatively recruits Aβ (10 – 25 µA), Aδ (30 – 70 µA) and C- (100 to 600 µA) fibers (Nakatsuka *et al*., 2000; Daniele & MacDermott, 2009). DRS-evoked EPSCs in laminae II neurons were identified using the following criteria: 1) constant response latency (i.e., response “jitter” < 200 μ) that did not change with increasing intensity of dorsal root stimulation; and 2) EPSCs that follow stimulation frequency ≤ 10 Hz for Aδ and ≤ 2 Hz for C-fibers with constant latency.

### Voltage-activated currents

SDH neurons display voltage-activated fast and slow A-type potassium currents (I_A_) (Ruscheweyh *et al*., 2004; Yasaka *et al*., 2010; Smith *et al*., 2015). To initiate our study of I_A_ in Y1eGFP neurons, we first segregated voltage-activated fast and slow transient outward currents (TOCs) based on their decay time constants in normal aCSF. To determine voltage-dependent activation, the cell membrane was voltage-clamped at -60 mV, and after a hyperpolarizing step to -100 mV for 250 ms, depolarizing voltage steps were applied at 5 s intervals in +5 mV increments from -80 mV to -40 mV (duration = 1 s). To determine voltage-dependent inactivation, the cell was voltage-clamped at -60 mV, and then prepulse voltage steps ranging from -110 mV to -40 mV were applied at 5 s intervals in +5 mV increments (duration = 150 ms) followed by a depolarizing step to -40 mV (duration = 1 s). The decay phase of the TOC (from time of peak to 250 ms) was fitted with an exponential equation [I = A·exp(-t/τ) + I_0_], where τ was the decay time constant, A the current amplitude and I_0_ is the initial current. Fast and slow A-currents were segregated based on the decay time constant. The above measurements of TOCs were refined to determine activation/inactivation curves of A-type currents. First, to isolate I_A_, 2 µM tetrodotoxin (TTX), 10 mM tetraethylammonium (TEA) and 200 µM Cd^2+^ were added to the circulating aCSF. Second, we increased the incremental step and extended the voltage range, while keeping the other parameters the same (Hu *et al*., 2003, 2006). To obtain activation curves, the maximum step was extended to +70 mV. To obtain inactivation curves, step to voltage was changed to +40 mV. For both activation and inactivation curves, 10 mV incremental steps were used (instead of 5 mV). In parallel, we also obtained baseline currents by repeating these activating and inactivating voltage steps following a 150 ms step to -10 mV. We offline subtracted this baseline current from the corresponding TOC to obtain I_A_ (Hu *et al*., 2003, 2006). Activation and inactivation curves were obtained from peak values of I_A_ for each voltage step. The voltage dependence of activation and inactivation of I_A_ was fitted with the Boltzmann function as follows. For activation, peak currents were converted to conductance (G) by the formula G = I/(V_m_ -V_rev_), where I is the current, V_m_ is the membrane voltage of depolarization pulses, and V_rev_ is the calculated potassium reversal potential (-88.3 mV). The function G/G_max_ = 1/(1 + exp[(V1/2 - V)/k]) was used with the normalized conductance, where G_max_ is the maximal conductance obtained with a depolarizing pulse to +70 mV, V_1/2_ is the half-maximal voltage (V), and k is the slope factor. For inactivation, I/I_max_ + 1/(1 + exp[(V_1/2_ - V)/k]) was used, where I_max_ is the maximal current obtained with a -110 mV prepulse. To determine current density at +40 pA, we divided the peak value of I_A_ (at the voltage step to +40 mV, from inactivation recordings) by the respective cell capacitance. <u>Transient inward currents</u>: Some neurons (before pharmacological isolation of I_A_) exhibited transient inward currents instead of TOCs. To further study these, we applied depolarizing current steps from -100 mV to -50 mV at 5 s intervals in +5 mV increments for 250 ms. These analyses were performed in the absence of cation channel blockers. <u>Hyperpolarization-activated currents</u>: Similarly, some neurons bathing in aCSF exhibited slow hyperpolarization-activated inward currents. To further study these, we applied hyperpolarizing current steps from -50 mV to -100 mV at 5 s intervals in +10 mV increments for 1 s duration.

### Pharmacology

Drugs were bath-applied using a valve-controlled perfusion system. To test the functional responsiveness of Y1-expressing neurons, we rapidly applied 20 µM NPY (Bachem, Switzerland) or the Y1 selective agonist [Leu^31^,Pro^34^]-NPY (Anaspec, USA) by pressure ejection with a pipette positioned close (∼50 µm) to the soma and evaluated inwardly rectifying K^+^ currents. Before and during the application of drug, voltage ramps (from -50 to -130 mV) were applied at a rate of 0.2 mV/ms. The NPY-sensitive component of the current was determined by offline subtraction of currents obtained during drug application from those obtained before drug application. 5s after NPY washout, the voltage ramp protocol was repeated to determine the recovered current. NPY-mediated conductance (*g*) was calculated as: *g = I/(V_m_ - E_rev_)* at two diL ent membrane potentials (*V_m_*) that were equidistant (25 mV) from the reversal potential (*E_rev_*). To estimate the degree of inward rectification, the ratio of these two conductance was calculated as: *g(E_rev_+25)/g(E_rev_-25)*. *E_rev_* is the potential (*V*) at which *I* = 0.

To determine whether the transient outward current evoked by the step potential from - 100 mV to +40 mV are I_A_, we blocked the currents by application of 0.5 and 5 mM 4- aminopyridine (4-AP) in the presence of 2 µM TTX, 10 mM TEA and 200 µM Cd^2+^. To determine whether the transient inward current evoked by a depolarizing step from -100 mV to -50 mV are low voltage-activated T-type calcium currents (I_Ca,T_), we blocked the currents by application of 100 µM Ni^2+^ (Yasaka *et al*., 2010; Smith *et al*., 2015). To determine whether the slow inward currents elicited by current steps from -50 mV to -100 mV are hyperpolarization-activated inward currents (I_h_), we blocked the currents by application of 2 mM Cs^+^ (Hughes *et al*., 2012; Smith *et al*., 2015).

### Statistics

Data management and simple statistical tasks were carried out with Clamp fit 10.3 (Molecular Devices, USA). Further statistical analysis was performed using Sigma Plot 12.0 (Systat, USA), MATLAB 2016b (The MathWorks, USA), and GraphPad Prizm 6.0 (GraphPad Software, USA). Analysis of variance (ANOVA) (for drug and A-current interaction), t-test, χ test, Fisher’s exact test, Mann-Whitney rank sum test and K-means clustering, were used for statistical comparison (see Statistical Summary Table for detail). Single comparisons between groups were made by the unpaired *t* test. If a set of data did not have a normal distribution, confirmed by the Shapiro–Wilk test, the Rank Sum test was used for the comparisons between groups. Significance was set at p<0.05. The results are expressed as mean ± standard deviation (SD).

## RESULTS

### Expression of Y1-mRNA in the Y1eGFP mouse dorsal horn

Using immunohistochemistry in rat spinal cord, at least seven distinct populations of Y1R-expressing interneurons have been identified, the most abundant of which are densely packed within the superficial dorsal horn, particularly in outer lamina II (Brumovsky *et al*., 2006, 2007; Hökfelt *et al*., 2007). To determine whether this pattern is reproduced in the Y1eGFP mouse, we first examined the relationship between GFP immunohistochemistry and Y1-mRNA expression in perfusion-fixed tissue. As illustrated in **Figure 2A**, we found that GFP labelled profiles were distributed throughout the laminae I-III, with highest density in outer lamina II as predicted. We did not find GFP labelled profiles in deeper dorsal horn or ventral horn. As shown in Table 1, quantitative analysis revealed that the great majority (86%) of GFP labelled profiles throughout laminae I-III also expressed Y1 mRNA (Figures 2A-2E). Similar results were found when segregating the analysis to laminae I-II (88%) and lamina III (81%). The colocalization of Y1-mRNA and Y1-eGFP is greater in lamina I-II than in lamina III (see also table 1). Together, these data indicate that GFP labels Y1R-expressing neurons with high fidelity.

**Figure 2.**
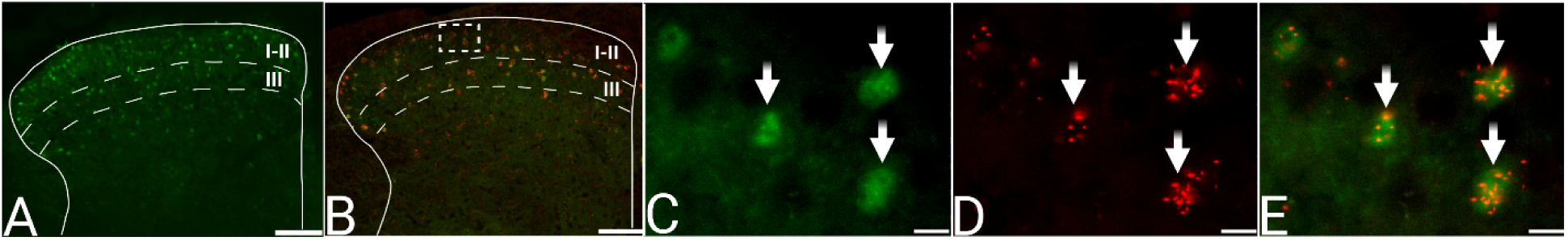
Expression of Y1-mRNA in the Y1eGFP mouse dorsal horn. (**A**) Distribution of green fluorescent protein (GFP) in the dorsal horn of a representative Y1eGFP mouse. Dotted lines represent boundaries of the dorsal horn laminae I-II and III(Peirs *et al*., 2020)(Peirs *et al*., 2020)(Peirs *et al*., 2020)(Peirs *et al*., 2020)(Peirs *et al*., 2020)(Peirs *et al*., 2020)(Peirs *et al*., 2020)(Peirs *et al*., 2020)(Peirs *et al*., 2020)(Peirs *et al*., 2020)(Peirs *et al*., 2020)(Peirs *et al*., 2020)(Peirs *et al*., 2020)(Peirs *et al*., 2020)(Peirs *et al*., 2020)(Peirs *et al*., 2020)(Peirs *et al*., 2020)(Peirs *et al*., 2020)(Peirs *et al*., 2020)(Peirs *et al*., 2020)(Peirs *et al*., 2020)(Peirs *et al*., 2020)(Peirs *et al*., 2020)(Peirs *et al*., 2020)(Peirs *et al*., 2020)(Peirs *et al*., 2020)(Peirs *et al*., 2020)(Peirs *et al*., 2020). (**B**) Fluorescent *in situ* hybridization experiments confirmed Y1-mRNA (red) expression in the majority of Y1eGFP neurons (green) throughout laminae I-III. The box inset represents the area of the dorsal horn that is zoomed in 10-fold for panels C-E. (**C**) Y1eFGFP (green), (**D**) Y1-mRNA (red puncta) and (**E**) merged (co-localization) as denoted by the arrows. Scale bars: 100 µm (A - B) and 10 µm (C - E).

**Table 1:** Y1-mRNA and Y1eGFP co-expression in laminae I-III of L3-L5 spinal cord. Y1-mRNA was not detected in lamina IV. Shown are the total number of DAPI-positive profiles double labelled with either Y1eGFP or Y1-mRNA, or triple labelled with both Y1eGFP and Y1-mRNA. Also shown are the percentage of Y1-mRNA positive cells that co-expressed Y1eGFP, and the percentage of Y1eGFP positive cells that co-expressed Y1-mRNA. Values are mean ± SD, N=4 mice (4 sections averaged for each of the 4 mice).

| Laminae | Y1eGFP<br>(total) | Y1-mRNA<br>(total) | Y1eGFP +<br>Y1-mRNA | % Y1eGFP that<br>are Y1-mRNA | % Y1-mRNA that<br>are Y1eGFP |
| --- | --- | --- | --- | --- | --- |
| I-III | 30.4 ± 6.7 | 40.0 ± 7.2 | 26.1 ± 5.6 | 84.8 ± 7.6 | 64.1 ± 4.6 |
| I-II | 21.1 ± 4.4 | 27.5 ± 5.1 | 18.6 ± 3.9 | 88.2 ± 9.7 | 68.8 ± 3.1 |
| III | 9.3 ± 2.3 | 12.5 ± 2.1 | 7.5 ± 1.7 | 81.4 ± 5.6 | 59.3 ± 6.1 |

### Effect of Y1 agonists on the membrane potential of Y1eGFP neurons

To study the functional responsiveness of Y1eGFP neurons, we applied Y1R agonists to spinal cord slices. As previously described (Miyakawa *et al*., 2005; Melnick, 2012), bath administration of NPY or the Y1R-specific agonist [Leu^31^,Pro^34^]-NPY produced outward currents and membrane hyperpolarization in dorsal horn interneurons. As illustrated in **Figure 3A**, application (15 s) of NPY (20 µM) induced outward currents (ΔI > 5pA were considered as positive response) in 8 (4 animals) of 34 Y1eGFP neurons. Application of aCSF vehicle never changed holding current (n = 4 neurons; 3 animals), ruling out an effect of the pressure wave (Fig 3B). As illustrated in Figure 3C, Leu^31^,Pro^34^]-NPY (20 µM) induced outward currents following agonist application in 4 (3 animals) of 11 neurons. In two of the four responding neurons, we applied the agonist for a longer duration (170 s). In these cases the outward current gradually subsided at the end of the agonist application, perhaps due to receptor desensitization (Rajagopal & Shenoy, 2018). As illustrated in Figure 3D, the peak of the outward current produced by NPY was similar to current peaks by [Leu^31^,Pro^34^]-NPY, approximately 14 pA. Under current clamp conditions, Figure 3E-F illustrate that application of NPY (20 µM) induced a hyperpolarization of 10 ± 1 mV in 3 Y1eGFP neurons recorded from 3 different animals.

**Figure 3.**
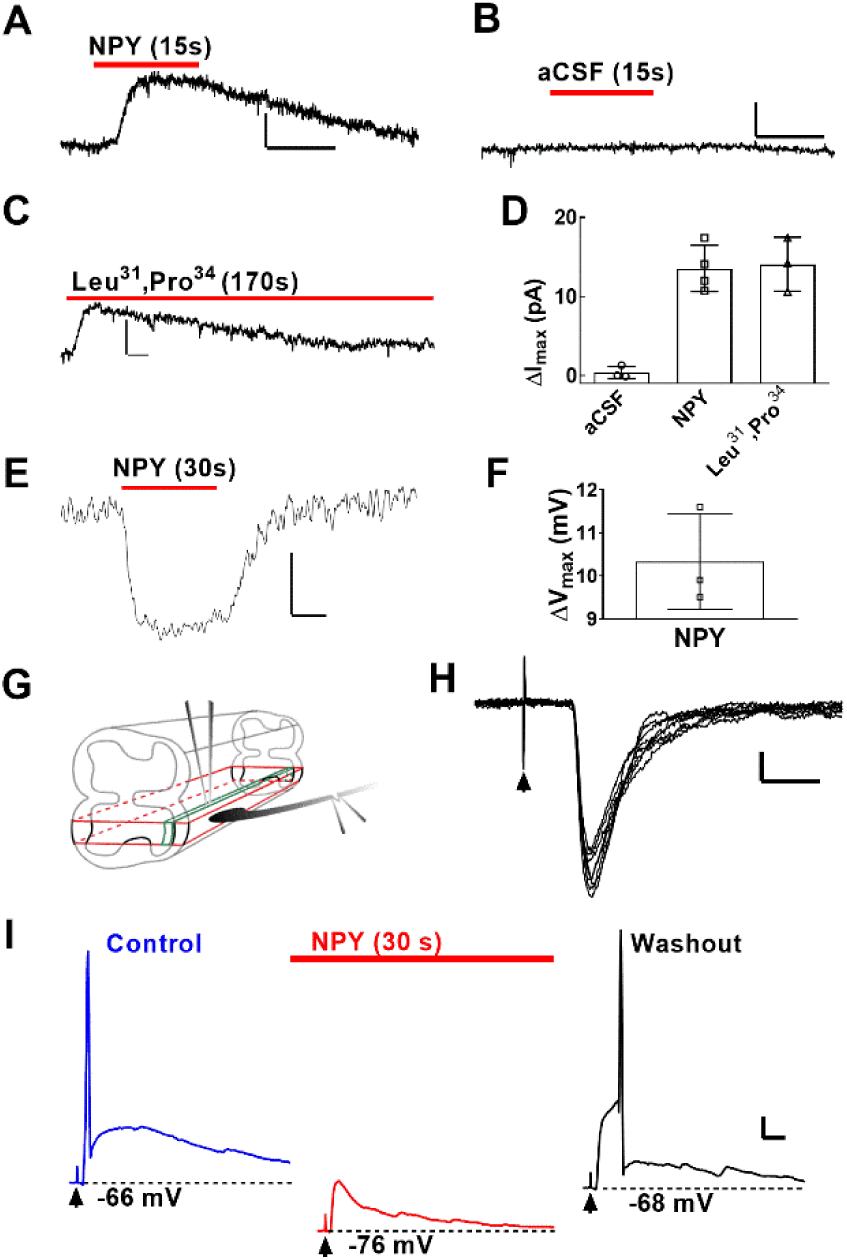
Membrane properties and dorsal root stimulus-evoked EPSCs in dorsal horn Y1eGFP interneurons after application of Y1R-selective agonists. Representative traces of outward currents in Y1eGFP neurons held at V_h_ = -60 mV following puff application of (**A**) 20 µM NPY, (**B**) aCSF or (**C**) 20 µM [Leu^31^,Pro^34^]-NPY. horizontal bar = 10 s, vertical bar = 10 pA. (**D**) Peak current amplitude within 10 s of application of aCSF (N=3 mice, with 2 neurons averaged within 1 of the mice), NPY (N=4 mice; 8 neurons responded out of a total of 34 Y1eGFP neurons given drug), or [Leu^31^,Pro^34^]-NPY (N=3 mice, 4 neurons responded out of a total of 11 neurons given drug). (**E)** Representative trace and (**F)** mean change in membrane potential (hyperpolarization from RMP) of Y1eGFP neurons following rapid application of 20 µM NPY (N=3 mice, one neuron per mouse). horizontal bar = 10 s, vertical bar = 5 mV. (**G**) Schematic of a parasagittal slice from the lumbar spinal cord with dorsal root attached. Dorsal root stimulation (DRS) was facilitated by glass pipette with electrodes and attached to the dorsal root by suction as illustrated. (**H)** Monosynaptic EPSCs following DRS at A-δ recruiting strengths (arrowhead). horizontal bar = 20 ms, vertical bar = 20 pA. **(I)** Current clamp recordings taken before (blue line, left), during (red line, middle), and 120 s after (black line, right) application of NPY in the setting of DRS at A-δ recruiting strength. Before NPY, DRS evoked a single action potential (blue line, left). Puff application of NPY hyperpolarized the cell and prevented DRS- evoked action potential firing (red line, middle). 120 s after NPY washout, the cell nearly returned to resting membrane potential, and DRS again evoked an action potential (black line, right). horizontal bar = 50 ms, vertical bar = 5 mV. Arrowheads indicate time of DRS. Values in D and F represent mean ± SD; each dot represents average value of one mouse.

### Dorsal root stimulation evokes action potential firing in Y1eGFP neurons: inhibition by NPY

Figure 3G illustrates our spinal cord slice preparation. The voltage clamp recordings of Figure 3H demonstrates for the first time in Y1eGFP neurons that DRS evoked monosynaptic (75%) and polysynaptic (25%) post-synaptic currents at A-δ (n=4) recruiting strengths (30-70 μA), as well as monosynaptic (37%) and polysynaptic (63%) post-synaptic currents at C-fiber (n=8) recruiting strengths (70-1000 μA). The current clamp recording in Figure 3I illustrate that DRS at A-δ recruiting strength evoked an action potential.

Aδ- and C-fibers are essential for the transmission of noxious somatosensation to the CNS (Todd & Koerber, 2006; Yasaka *et al*., 2010), but Miyakawa et al [2005] reported that NPY did not change either Aδ- or C-afferent fiber-mediated eEPSPs in unidentified rat dorsal horn neurons. To re-evaluate this question in identified Y1R-expressing neurons, we determined whether NPY changes DRS-evoked activity in Y1eGFP neurons. NPY (20 µM) reversibly hyperpolarized the membrane by ∼10 mV and abolished DRS-evoked action potential firing (Figure 3I). These data indicate that NPY inhibits the responsiveness of Y1eGFP neurons to primary afferent input.

### Effect of Y1R agonists on inwardly rectifying K^+^ currents in Y1eGFP neurons

In unidentified dorsal horn neurons, NPY induces outward currents that can be blocked with either barium ions or cesium ions + TEA, consistent with activation of K^+^ channels (Miyakawa *et al*., 2005; Melnick, 2012). The NPY current was also blocked with GDP-β-S (Miyakawa *et al*., 2005; Melnick, 2012), and current-voltage curves revealed that NPY generated inward rectification in some neurons (Moran *et al*., 2004; Miyakawa *et al*., 2005), suggesting an NPY-mediated activation of Gi-protein coupled inwardly rectifying K^+^ (GIRK) channels. To provide a more quantitative analysis of GIRK channel activity in identified Y1R- expressing neurons under voltage clamp conditions, we obtained currents elicited by voltage ramps delivered to spinal cord slices from Y1eGFP mice, and then generated I-V curves before (control), during, and after the rapid application of NPY (**Figure 4A**) or [Leu^31^,Pro^34^]-NPY. We then subtracted control currents from currents recorded during NPY or [Leu^31^,Pro^34^]-NPY application (Figure 4B-C).

**Figure 4.**
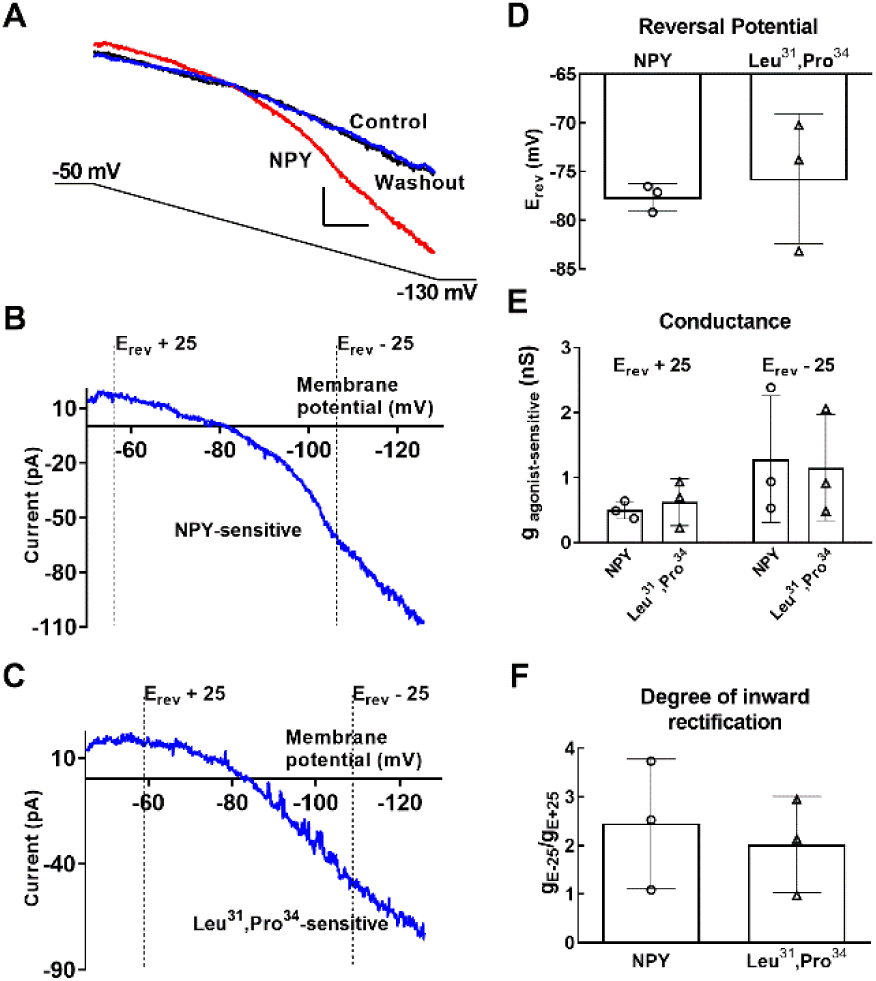
Y1 receptor agonists induce inwardly rectifying currents in Y1eGFP neurons. **(A)** Voltage ramp-induced currents before (blue line, control), during (red line, NPY) and immediately after (black line, washout) rapid application of 20 µM NPY. The voltage-ramp protocol is shown at bottom (gray line). (**B**) NPY-sensitive current calculated with off-line subtraction of Control from NPY. (**C**) [Leu^31^,Pro^34^]-NPY-sensitive current calculated with off-line subtraction of Control from [Leu^31^,Pro^34^]-NPY. (**D**) Mean reversal potentials [*E_rev_*] for current induced by 20 µM NPY (N=3, 5 neurons) or 20 µM [Leu^31^,Pro^34^]-NPY (N=3, one neuron per mouse). (**E**) Conductance (*g*) measured ±25 mV from *E_rev_* (N=3, one neuron per mouse). (**F**) Degree of inward rectification (*g_(Erev + 25) /_ g_(Erev – 25)_*) of NPY- or [Leu^31^,Pro^34^]-NPY-sensitive currents (N=3, one neuron per mouse). Values in D, E and F represent mean ± SD; each dot represents average value of one mouse.

The reversal potential (*E_rev_*) reflects the Nernst equilibrium potential for K^+^ when adjusted for junction potential. As illustrated in Figure 4D, *E_rev_* for NPY- and [Leu^31^,Pro^34^]-NPY-induced currents was -77 ± 4 mV (n=5 neurons from 3 animals) and -76 ± 7 mV (n=3 neurons from 3 animals), respectively. Further inspection of Figures 4B-C reveals that the slope of the I-V curve was greater at potentials negative to E_rev_ compared to potentials positive to E_rev_. Subsequent analyses (Figure 4E) indicate inward rectification of NPY- and [Leu^31^,Pro^34^]-NPY-mediated K^+^ currents (Figure 4F). Taken together, the above data indicate that Y1R agonists open GIRK channels to hyperpolarize Y1-INs and thus block action potential discharge, leading to the occlusion of spinal transmission of nociceptive signals from the central terminal of primary afferent neurons.

### Intrinsic membrane properties of Y1eGFP neurons

A summary of intrinsic membrane properties of Y1eGFP neurons and randomly sampled neurons in SDH is illustrated in **Table 2**. We found that passive membrane properties, including membrane capacitance, input resistance and resting membrane potential, were similar between Y1eGFP and randomly sampled neurons. Also, active membrane properties, including action potential amplitude and afterhyperpolarization amplitude, were similar between Y1eGFP and randomly sampled neurons. However, action potential threshold was less-negative (more depolarized, p=0.0007, Statistical Summary Table P1) and action potential width was wider in Y1eGFP neurons as compared to randomly sampled neurons (p=0.009, Statistical Summary Table P1).

**Table 2:** Passive membrane properties [membrane capacitance, input resistance, and resting membrane potential (RMP)] and active membrane properties [action potential (AP) threshold, width, height, and afterhyperpolarization (AHP)] of randomly sampled (Random) and Y1eGFP SDH neurons. Data were collected from 14 WT mice (for recording of randomly sampled neurons) and 27 Y1eGFP mice. 1-5 neurons averaged for each mouse. Values represent mean ± SD. Numbers of animals are indicated in parentheses. **p<0.01 or ***p<0.001, Student’s t-test, between respective injecting strengths, Random vs. Y1eGFP.

|  | Random | Y1eGFP |
| --- | --- | --- |
| Membrane capacitance (pF) | 26 ± 6 (10) | 29 ± 7 (22) |
| Input resistance (MΩ) | 679 ± 380 (11) | 702 ± 201 (22) |
| RMP, mV | -65 ± 6 (10) | -61 ± 6 (22) |
| AP threshold (mV) | -39.2 ± 4.0 5(10) | -32.3 ± 3.0 (22)*** |
| AP width at base (ms) | 1.9 ± 0.3 (11) | 2.3 ± 0.3 (22)** |
| AP height from base (mV) | 82 ± 11 (11) | 84 ± 7 (22) |
| AHP amplitude (mV) | 20.5 ± 4.1 (11) | 20.2 ± 3.2 (22) |

### The action potential discharge patterns of Y1eGFP neurons include a unique, adapting form of delayed short latency firing

The firing patterns of action potentials can be used to segregate different neuronal populations in superficial dorsal horn (Todd, 2015, 2017). Depolarizing current injection typically evokes several distinct patterns that are often but not exclusively referred to as initial bursting firing (IBF), phasic firing (PF), tonic firing (TF), and delayed firing (DF) (Prescott & De Koninck, 2002; Ruscheweyh & Sandkühler, 2002; Ruscheweyh *et al*., 2004; Balasubramanyan *et al*., 2006; Schoffnegger *et al*., 2006; Yasaka *et al*., 2010; Punnakkal *et al*., 2014; Smith *et al*., 2015; Stebbing *et al*., 2016). Current injection is typically injected while the cell is held at resting membrane potential (Smith *et al*., 2015); however, detection of the delayed firing pattern often requires current to be injected from hyperpolarized holding potentials (Schoffnegger *et al*., 2006). Therefore, to determine the effect of membrane potential on evoked firing patterns, we injected depolarizing current not only at a holding potential at or near RMP (range: -65 to -80 mV), but also from a depolarized holding potential (-50 to -65 mV) and from a hyperpolarized holding potential (range: -80 to -95 mV).

We identified five main firing patterns with the following characteristics: <u>Delayed, Short Latency Firing (DSLF)</u> neurons were defined as those exhibiting a slow depolarizing delay of 60- 180 ms prior to firing the first action potential. <u>Delayed, Long Latency Firing (DLLF)</u> neurons were defined as those that exhibited a delay in firing of >180 ms (range, 180-850 ms). <u>Initial Burst Firing (IBF)</u> neurons were defined as those exhibiting a short burst of two to four high frequency (∼90 Hz) action potentials immediately after onset of current injection. <u>Tonic Firing (TF)</u> neurons exhibited continuous high frequency (≥40 Hz) firing immediately after onset of current injection. <u>Phasic Firing (PF)</u> neurons typically exhibited several action potentials immediately after the onset of current injection but terminated before the end of the current pulse.

In randomly sampled superficial neurons held at depolarized, resting, and hyperpolarized potentials (**Figure 5**), current injection evoked DSLF, DLLF, IBF or TF patterns. DSLF neurons typically exhibited irregular firing at weaker current injections of 40-80 pA, but more regular, non-adapting firing at higher injection strengths. At the onset of injection, most DLLF neurons exhibited a characteristic hyperpolarizing “notch” on the voltage shoulder, followed by a slow depolarizing ramp delay (Luther *et al*., 2000). Several DLLF neurons exhibited properties of gap firing neurons (Ruscheweyh *et al*., 2004; Schoffnegger *et al*., 2006), in that they spiked at the onset of current injection (mostly at higher depolarizing currents) and then, following a long delay, exhibited regularly spaced action potential firing. IBF neurons exhibited high frequency bursts of action potentials at middle current injection strengths before switching to adaptive firing at higher strengths when evoked from hyperpolarized holding potentials, as was the case for lamina I projection neurons in rats (Ruscheweyh *et al*., 2004). Furthermore, the firing frequency decreased at higher depolarizing steps and adapted. The firing frequency of TF neurons increased in proportion to current strength but did not change during a current injection step. In most TF neurons the firing did not cease even at the highest 300 pA step. PF neurons typically exhibited action potentials patterns like DSLF neurons but different in that they occurred immediately after the onset of current injection. Most action potentials were continuous while a few others adapted (frequency decreased) with time (Yasaka *et al*., 2010; Boakye *et al*., 2018). In most cases the firing ceased before reaching 150 pA.

**Figure 5.**
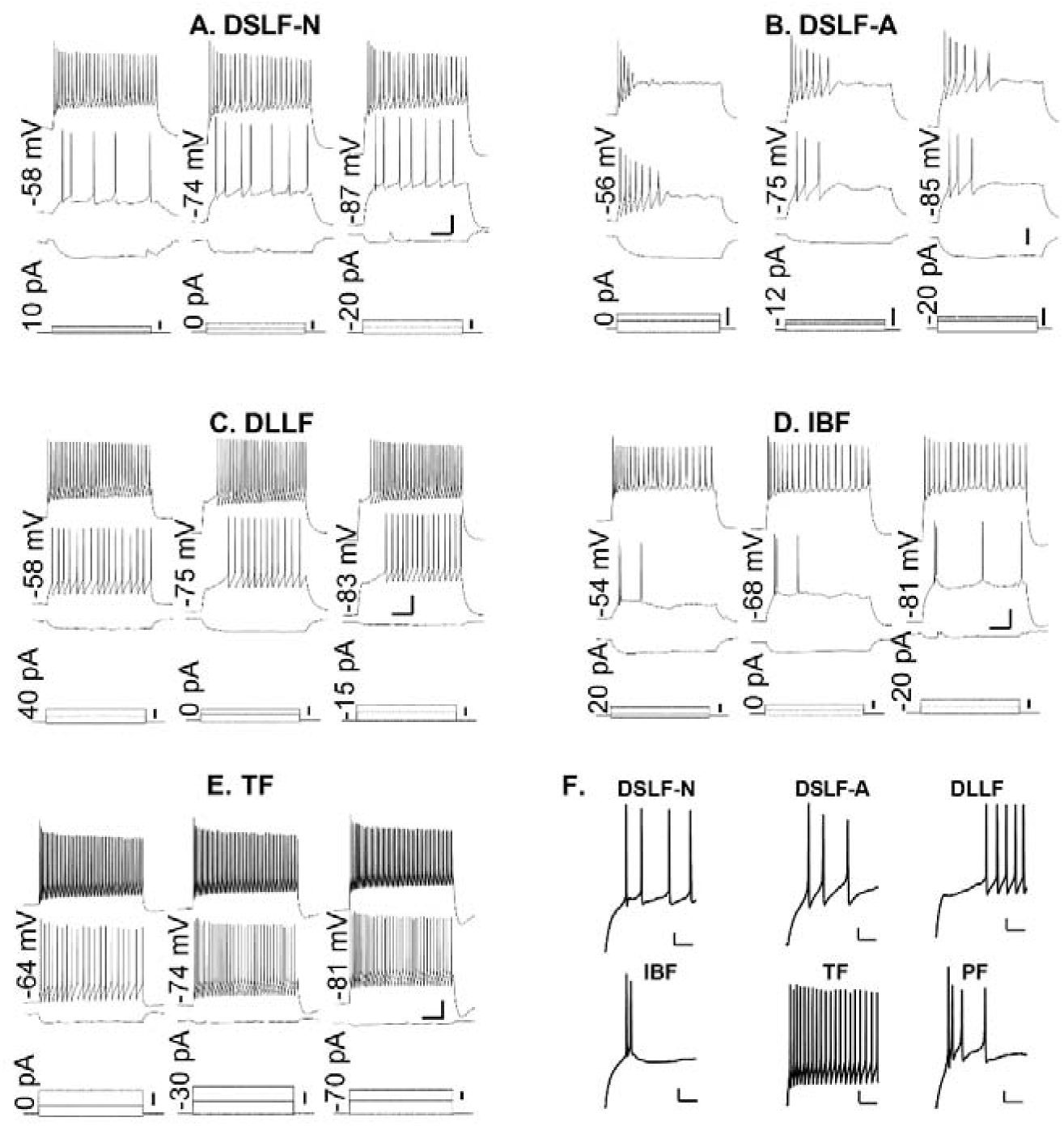
The action potential discharge patterns of Y1eGFP neurons include a unique, adapting form of delayed short latency firing. The predominant action potential firing patterns recorded after steady-state current injection (multiple steps of 1 s each) from randomly sampled (RS) and Y1eGFP (Y1) neurons: (**A-E**) DSLF-N (RS): delayed short latency – non-adapting, DSLF-A (Y1): delayed short latency – adapting, DLLF (RS): delayed long latency, IBF (RS): initial burst and TF (RS): tonic action potential firing in randomly sampled and Y1eGFP neurons. Action potential firing patterns were elicited from three conditioning membrane potentials represented within the panel for each FP: depolarizing (left figures), resting (middle figures) and hyperpolarizing (right figures), each shown for three current injection strengths as represented at bottom. The three current injection strengths were chosen to produce: hyperpolarization (bottom), reliable firing (middle), and maximal firing of APs (top). The values on the left side of each set of (three) firing patterns indicate the conditioning membrane potential that were obtained by the current injection magnitudes shown on the left of the respective CIs. Scale bars – horizontal = 200 ms (firing patterns and current steps), vertical = 20 mV (firing patterns) or 100 pA (current steps). (**F**): Enlarged images of each firing type, elicited from hyperpolarized conditioning membrane potentials with LCI. A representative example of a phasic firing (PF) neuron from a Y1eGFP slice indicates a firing pattern that is similar to DSLF-A, but without the delay. Scale bars – horizontal = 100 ms, vertical = 10 mV.

The firing patterns of Y1eGFP neurons were notably different from randomly sampled neurons. A striking difference was the relatively few Y1eGFP neurons that exhibited the TF firing pattern (2% of Y1eGFP vs 10% of randomly sampled). Another striking difference is that almost all Y1eGFP DSLF neurons exhibited adaptation (decreasing firing frequency over time illustrated as delayed short latency – adapting (DSLF-A in Fig 5). Thus, DSLF neurons can be segregated into two sub-categories: Adapting (DSLF-A) that are characteristic of Y1eGFP neurons; and non-adapting (DSLF-N) that are characteristic of randomly sampled neurons. Furthermore, PF neurons exhibited similar firing patterns to DSLF neurons but without a delay (Fig. 5F).

The firing properties of DLLF and IBF neurons were similar when recorded from the randomly sampled or Y1eGFP populations. Also, the neuronal membrane capacitance, an indirect measure of cell body surface area, was significantly less for both randomly sampled and Y1eGFP-positive IBF neurons compared to non-IBF neurons (Random: 19±7 pF vs 32±9 pF, p=0.0005; Y1eGFP: 24±8 pF vs 31±9 pF, p=0.03).

We evaluated the firing patterns of Y1-eGFP neurons that responded to Y1 receptor agonists with those that did not. The percentage of DSLF, DLLF, IB and PF neurons that responded to Y1 agonists were 57%, 36%, 0% and 7%, while the percentage that did not respond were 47%, 24%, 19% and 9%, respectively.

To begin to identify the primary afferents that project to each type of Y1eGFP neuron, we first determined firing pattern in current clamp mode, and then switched to voltage clamp mode to evaluate DRS-evoked EPSPs. In DSLF neurons, we observed eEPSCs with a frequency of 71% (5/7) and 29% (2/7) at C- and Aδ recruiting strengths, respectively. In DLLF neurons, we observed eEPSCs with a frequency of 67% (2/3) and 33% (1/3) at C- and Aδ recruiting strengths, respectively. Both IBF neurons that we recorded from responded to C- but not Aδ recruiting strengths. This preliminary data set did not include PF or TF neurons.

### Firing patterns determine subpopulations of Y1eGFP neurons that are distinct from randomly sampled neurons

Previous studies have documented various firing patterns in superficial dorsal horn from either unidentified rat neurons (Prescott & De Koninck, 2002; Ruscheweyh & Sandkühler, 2002; Hantman *et al*., 2004; Ruscheweyh *et al*., 2004; Heinke *et al*., 2004) or GFP-labeled mouse neurons (Schoffnegger *et al*., 2006; Punnakkal *et al*., 2014; Smith *et al*., 2015). In **Figure 6**, we unveil the remarkably high impact of hyperpolarized conditions on the constellation of firing patterns in Y1eGFP neurons. When firing was evoked from resting membrane potential in randomly sampled neurons, we observed five major patterns in order of prevalence: DSLF > PF > DLLF/TF > IBF. When evoked in Y1eGFP neurons, we observed the following order of prevalence: PF >>> IBF > TF > DSLF/DLLF. When firing patterns were elicited from conditioning hyperpolarized potentials in the randomly sampled neurons, we noticed a small shift from PF and TF at RMP to DSLF and DLLF at hyperpolarized potentials. When, however, firing patterns were elicited from conditioning hyperpolarized potentials in the Y1eGFP population, we observed a major shift in prevalence from PF and TF to DSLF and DLLF.

**Figure 6.**
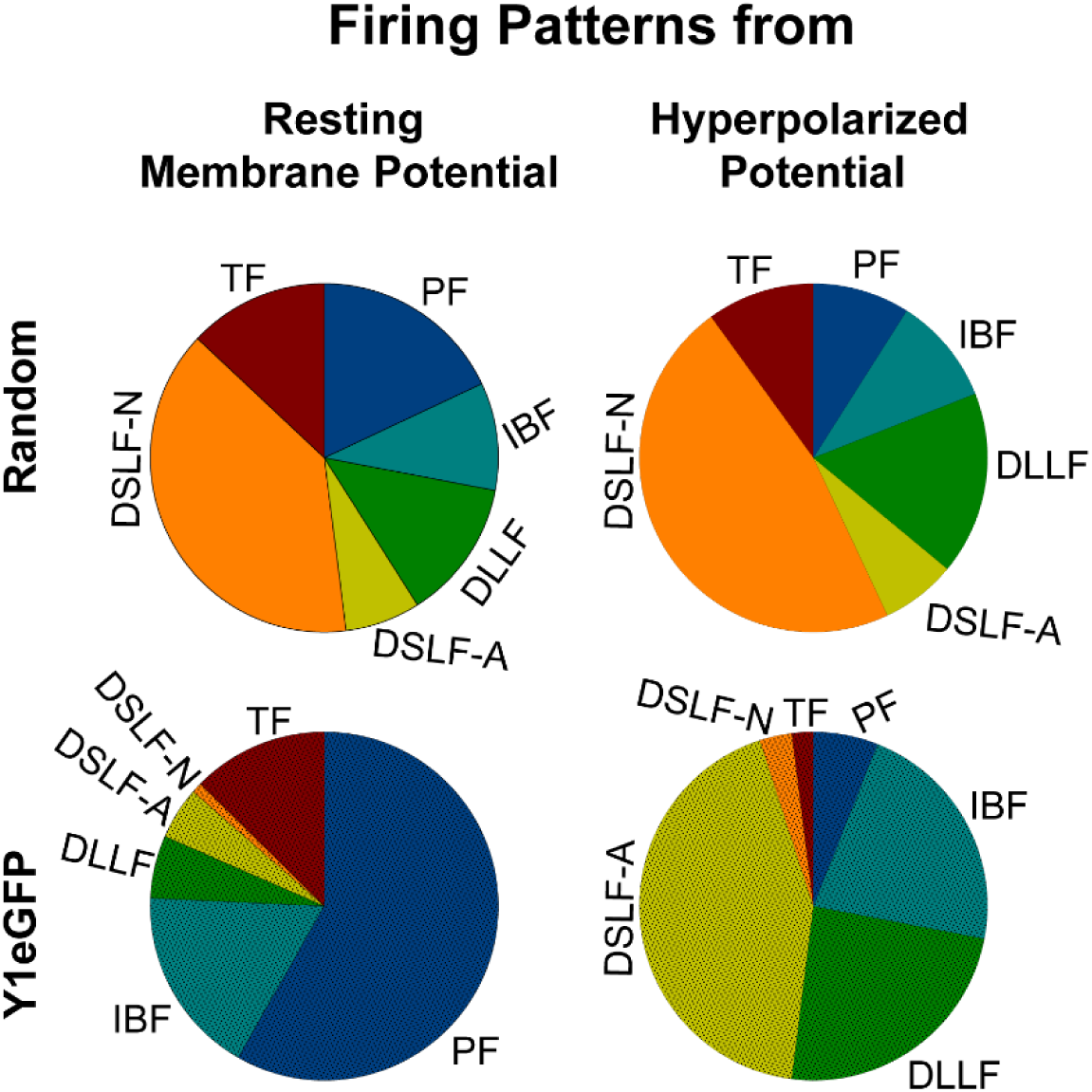
Firing patterns of Y1eGFP neurons are sensitive to resting membrane potential. Y1eGFP neurons undergo a dramatic shift from phasic and tonic firing patterns (evoked from at or near resting membrane potentials) to delayed short latency and long latency firing patterns (evoked from hyperpolarized potentials).

**Figure 7** illustrates the prevalence of the identified firing patterns elicited from conditioning hyperpolarized potentials in randomly sampled neurons from WT mice, and in Y1eGFP-negative and Y1eGFP-positive neurons from Y1eGFP mice. The firing patterns of Y1eGFP-negative neurons were essentially identical to the randomly sampled neurons (χ^2^ statistics; P=0.98) except for the TF type. Strikingly, the population distribution of firing patterns in Y1eGFP-positive neurons was significantly different from either randomly sampled or Y1eGFP-negative neurons (vs randomly sampled: χ^ 2^ statistics; P<0.001, vs Y1eGFP-negative: χ^2^ statistics; P<0.0001). We observed differences in percentages within individual firing patterns that contribute to overall significance. For example, in contrast to randomly sampled neurons (6/58 or 10%), TF was rarely observed in Y1eGFP-positive (2/120 or 2%) neurons (P=0.01, χ^2^ test). A much higher percentage or randomly sampled (27/58 or 47%) neurons exhibited DSLF-N than in Y1eGFP-positive (4/120 or 3%) neurons (P<0.0001, χ test). In contrast, DSLF-A was clearly prominent in Y1eGFP-positive neurons. Randomly sampled (4/58 or 7%) neurons exhibited DSLF-A in a much smaller percentage compared to Y1eGFP-positive (52/120 or 43%) neurons (P<0.0001, χ test). The frequency of DLLF and other subpopulations was similar between the two groups. The difference in IBF neuron percentage was not significant – 10% (6/58) and 22% (26/120). Overall, these differences indicate that Y1eGFP neurons comprise a distinct functional phenotype in the SDH in terms of action potential firing patterns.

**Figure 7.**
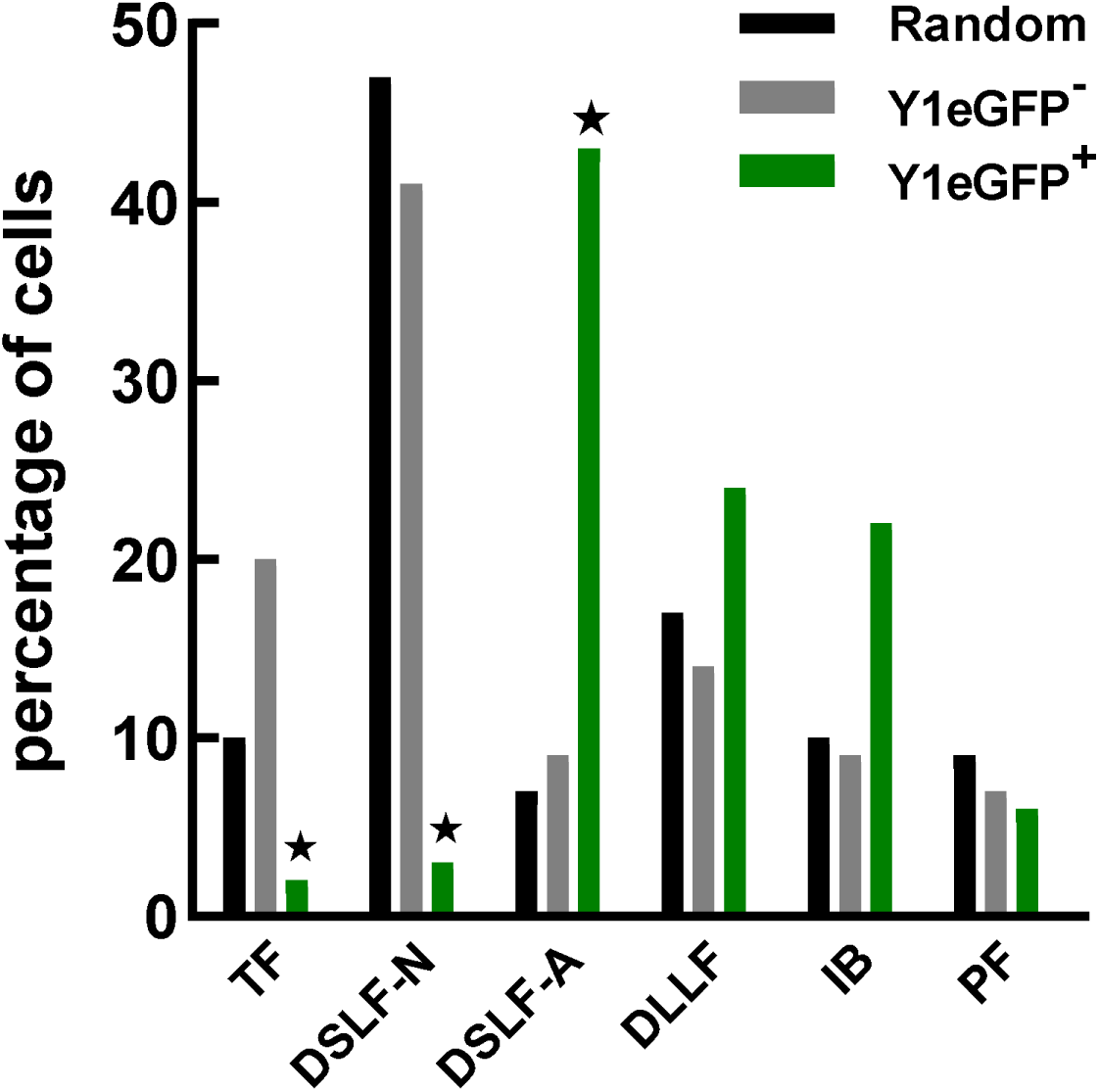
Firing patterns elicited from hyperpolarized conditions reveal Y1eGFP to label a distinct neuronal subpopulation. Incidence of tonic firing (TF), delayed short latency firing non-adapting (DSLF-N), delayed short latency firing adapting (DSLF-A), delayed long latency firing (DLLF), initial burst firing (IBF) or phasic firing (PF) patterns from hyperpolarized conditions in randomly sampled neurons (Random, n=58 from 14 wildtype mice), unlabeled neurons in Y1eGFP mice (Y1eGFP^-^, n=56 from 24 Y1eGFP mice) and Y1eGFP^+^ neurons in Y1eGFP mice (Y1eGFP^+^, n=120 neurons from 27 Y1eGFP mice). *p<0.05 Y1eGFP^+^ vs Random or eGFP^-^, Chi-square followed by Fisher’s exact test.

### Validation of distinct firing patterns in Y1eGFP neurons based upon k-means clustering

Classification of firing pattern, obtained from hyperpolarized conditions, were further refined by closer analysis of number of spikes, spike frequency, and spike frequency adaptation (Prescott & De Koninck, 2002). DSLF-A (randomly sampled) and DSLF-N (Y1eGFP) were not included for quantification. As summarized in **Tables 3-5**, quantitative analysis revealed some key differences in firing patterns: (a) shorter first spike latency to the onset of current injection in DSLF as compared to DLLF neurons (p<0.0001, Statistical Summary Table P2), regardless of whether they were non-adapting (randomly sampled) or adapting (Y1eGFP); (b) DSLF-A of Y1eGFP neurons exhibited fewer spikes (p<0.01, Statistical Summary Table P3), lower spike frequency (p<0.05, Statistical Summary Table P4), and greater adaptation (p=0.003, Statistical Summary Table P5) as compared to DSLF-N in randomly sampled neurons; c) IBF neurons had the highest spike frequency at small current injecting strengths; and d) TF neurons had the highest number of spikes and frequency at medium to large current injecting strengths. DLLF and TF neurons were quite similar in firing characteristics between Y1eGFP and randomly sampled neurons. More subtle differences between DSLF-N (randomly sampled) and DSLF-A (Y1eGFP) subpopulations were that DSLF-A (Y1eGFP) neurons exhibited fewer action potentials, more firing frequency adaptation, and shorter latency.

**Table 3:**
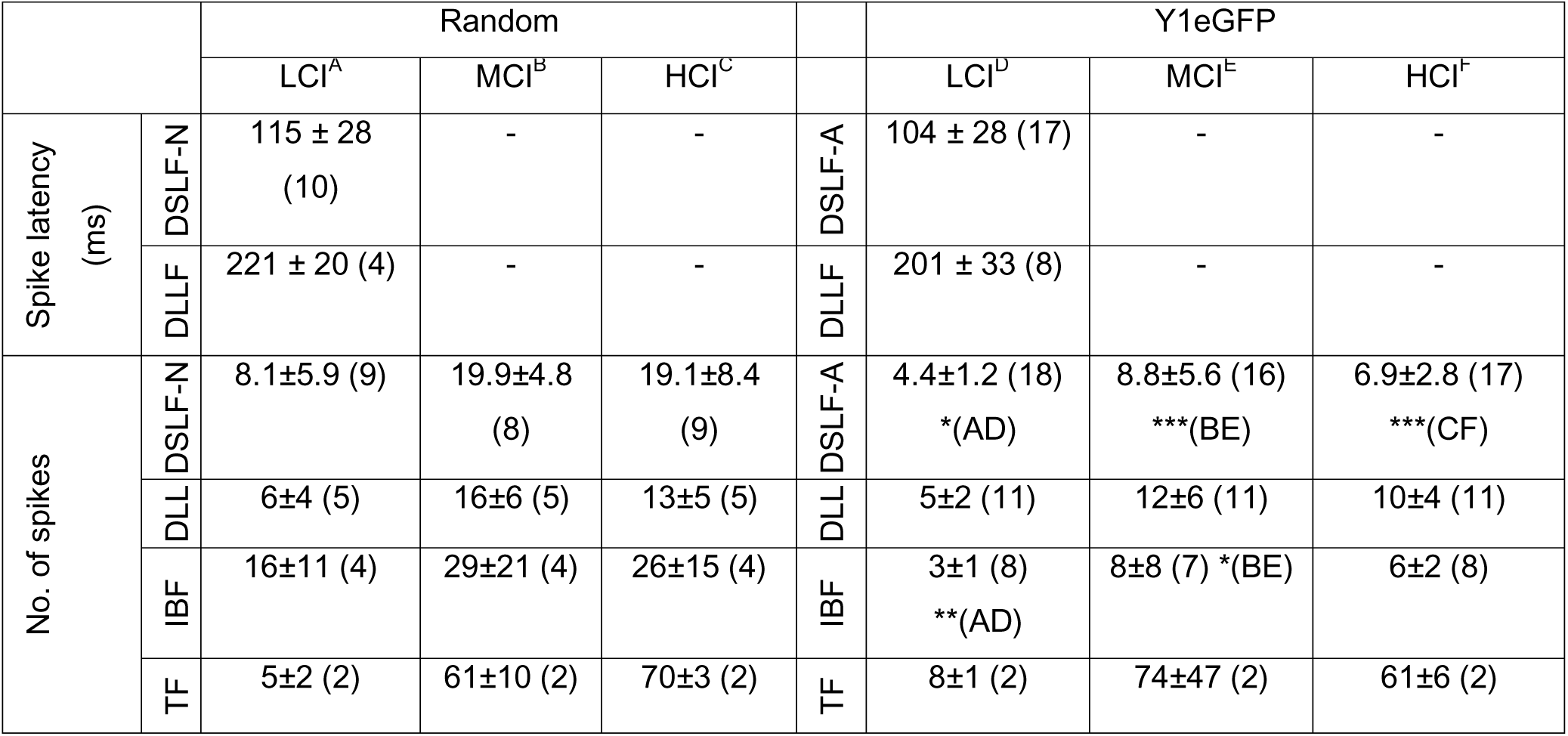
First spike latency in delayed firing neurons and number of spikes at different current injecting strengths from conditioning hyperpolarized potentials for randomly sampled (Random) and Y1eGFP SDH neurons. The latency to first spike was determined by injecting currents just above threshold (LCI). Numbers in brackets indicates number of animals. LCI is the first current injection step that elicited AP firing. HCI represents the magnitude of current injection generating the maximum number of APs. MCI represents the magnitude of current injection that was in between (the average of) LCI and HCI. DSLF stand for delayed short latency firing (N is non-adapting, A is adapting), DLLF for delayed long latency firing, IBF for initial burst firing and TF for tonic firing. 1-5 neurons averaged for each mouse. Values represent mean ± SD. *p<0.05, **p<0.01 or ***p<0.001, Student’s t-test, for respective injecting strengths, Random vs.

**Table 4:**
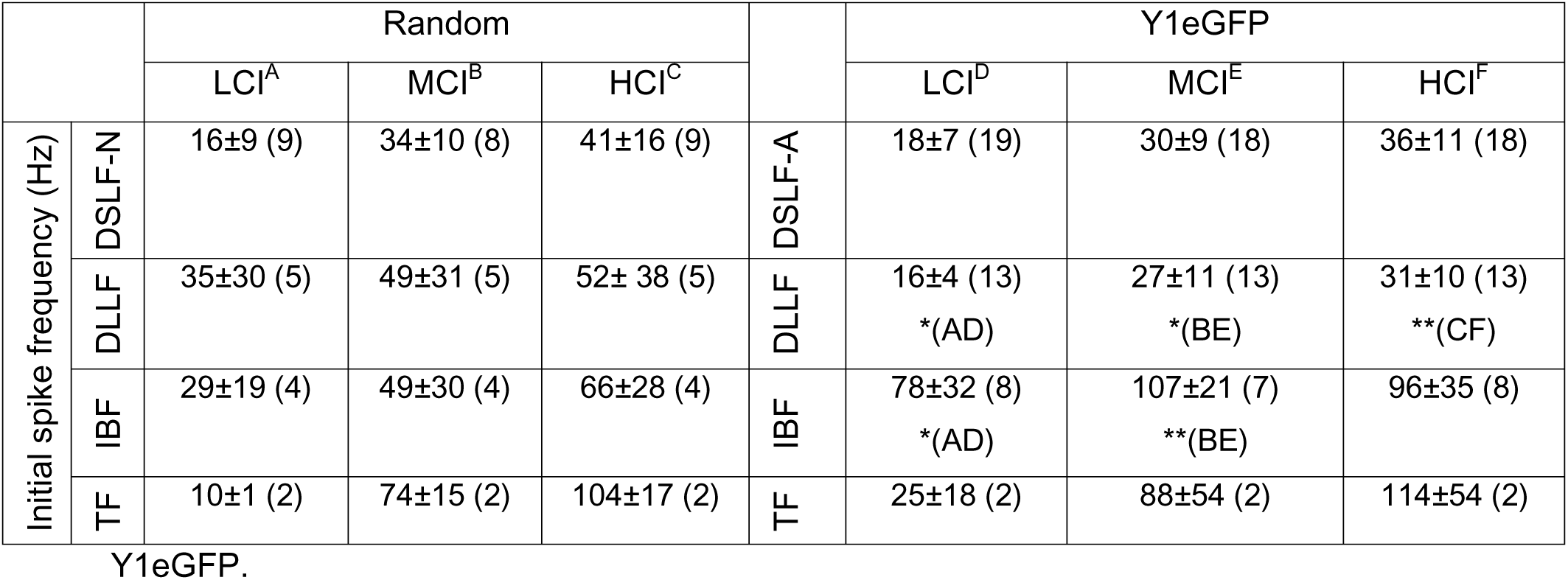
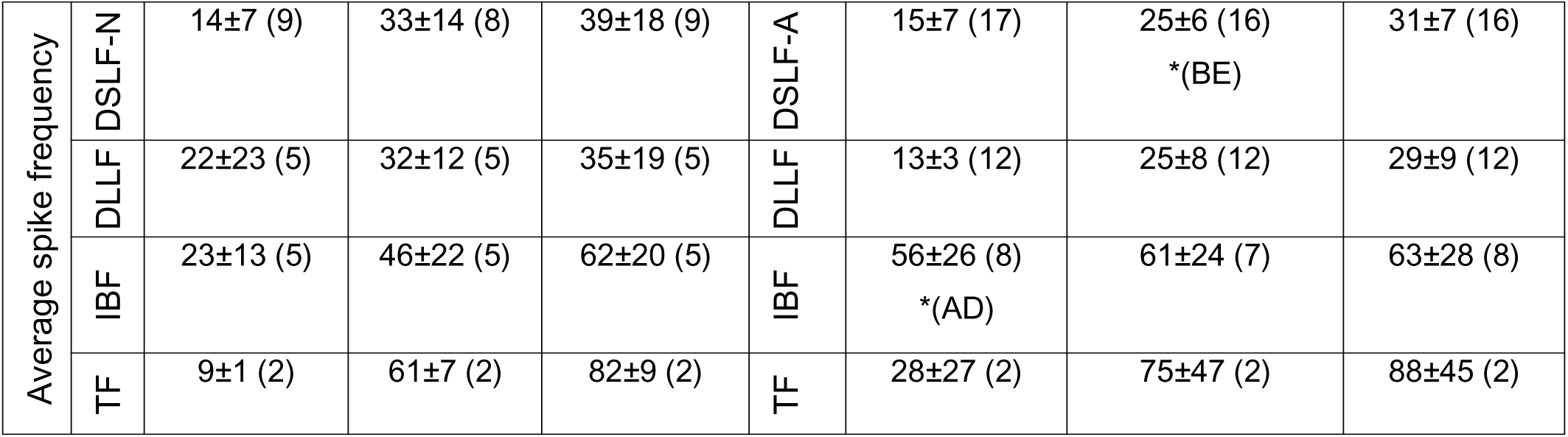
Characteristic firing frequencies (initial and average) at different current injection strengths from conditioning hyperpolarized potentials for randomly sampled (Random) and Y1eGFP SDH neurons. Numbers in brackets indicate number of animals. LCI, MCI and HCI denote Low, Medium, and High strengths of current injection. DSLF stand for delayed short latency firing (N is non-adapting, A is adapting), DLLF for delayed long latency firing, IBF for initial burst firing and TF for tonic firing. 1-5 neurons averaged for each mouse. Values represent mean ± SD. *p<0.05, **p<0.01 or ***p<0.001, Student’s t-test, for respective injecting strengths, Random vs. Y1eGFP.

**Table 5:** Characteristic frequency adaptation [initial {(*f*_2_ – *f*_1_)/ *f*_1_} and average of all {(*f_i_*_−1_ − *f_i_*)/*f_i_*}] at different current injection strengths from conditioning hyperpolarized potentials for randomly sampled (Random) and Y1eGFP SDH neurons. Number in brackets indicates number of animals. LCI, MCI and HCI denote Low, Medium and High strengths of current Injection. DSLF stand for delayed short latency firing (N is non-adapting, A is adapting), DLLF for delayed long latency firing, IBF for initial burst firing and TF for tonic firing. 1-5 neurons averaged for each mouse. Values represent mean ± SD. **\***p<0.05, **p<0.01 or ***p<0.001: Student’s t-test, for respective injecting strengths, randomly sampled vs. Y1eGFP.

|  |  | Random |  |  |  | Y1eGFP |  |  |
| --- | --- | --- | --- | --- | --- | --- | --- | --- |
|  |  | LCI<sup>A</sup> | MCI<sup>B</sup> | HCI<sup>C</sup> |  | LCI<sup>D</sup> | MCI<sup>E</sup> | HCI<sup>F</sup> |
| Initial frequency adaptation | DSL-F-N | -0.13±0.13 (9) | -0.01±0.19 (8) | -0.04±0.18 (9) | DSL-F-A | -0.21±0.14 (18) | -0.14±0.11 (17)<br>*(BE) | -0.10±0.09 (18) |
|  | DLL-F | -0.14±0.61 (5) | 0.07±0.23 (5) | -0.09±0.09 (5) | DLL-F | -0.15±0.16 (12) | 0.11±0.20 (12) | 0.09±0.16 (12)<br>*(CF) |
|  | IBF | 0.08±0.35 (4) | -0.16±0.27 (4) | -0.18±0.30 (4) | IBF | -0.47±0.09 (7)<br>**(AD) | -0.32±0.10 (7) | -0.25±0.35 (7) |
|  | TF | -0.09±0.01 (2) | 0.01±0.01 (2) | -0.04±0.02 (2) | TF | 0.33±0.12 (2)<br>*(AD) | 0.04±0.13 (2) | -0.01±0.05 (2) |
| average frequency | DSL-F-N | -0.05±0.13 (9) | -0.00±0.05 (8) | -0.03±0.04 (9) | DSL-F-A | -0.11±0.03 (15) | -0.07±0.05 (15)<br>**(BE) | -0.08±0.04 (17)<br>**(CF) |
|  | DLL | -0.04±0.13 (5) | -0.04±0.04 (5) | -0.05±0.03 (5) | DLL | -0.09±0.08 (12) | -0.03±0.03 (12) | -0.04±0.06 (12) |
|  | IBF | -0.01±0.11 (4) | 0.00±0.05 (4) | -0.02±0.03 (4) | IBF | -0.2±0.16(7)<br>*(AD) | -0.19±0.13 (7)<br>*(BE) | -0.12±0.08 (7)<br>*(CF) |
|  | TF | -0.42±0.43 (2) | -0.00±0.00 (2) | -0.00±0.00 (2) | TF | -0.15±0.17 (2) | -0.00±0.00 (2) | -0.01±0.01 (2) |

To validate the above unique properties of Y1eGFP neurons, we conducted a parameter-wise segregation of neurons into identifiable groups using k-means cluster analysis. We used two parameters – initial frequency and initial frequency adaptation for the analysis. As illustrated in Figure 8A, this analysis revealed that the randomly sampled neurons best clustered into four groups. Each sub-group exhibited a highly prevalent clustering of neurons within one of the following firing patterns: DSLF, DLLF, IBF and TF neurons (Table 6). On the other hand, Y1eGFP neurons best clustered into three groups: DSLF, DLLF and IBF neurons (Figure 8B). The analysis also revealed that the cluster center of DSLF (Y1eGFP) neurons tended towards greater adaptation as compared to the cluster center of DSLF (randomly sampled) neurons. In other words, the cluster center for DSLF (Y1eGFP) neurons was further below the computer-generated ‘zero line’ of initial frequency adaptation. TF Y1eGFP neurons did not cluster separately, probably because the incidence of occurrence was too low (2/120).

**Figure 8.**
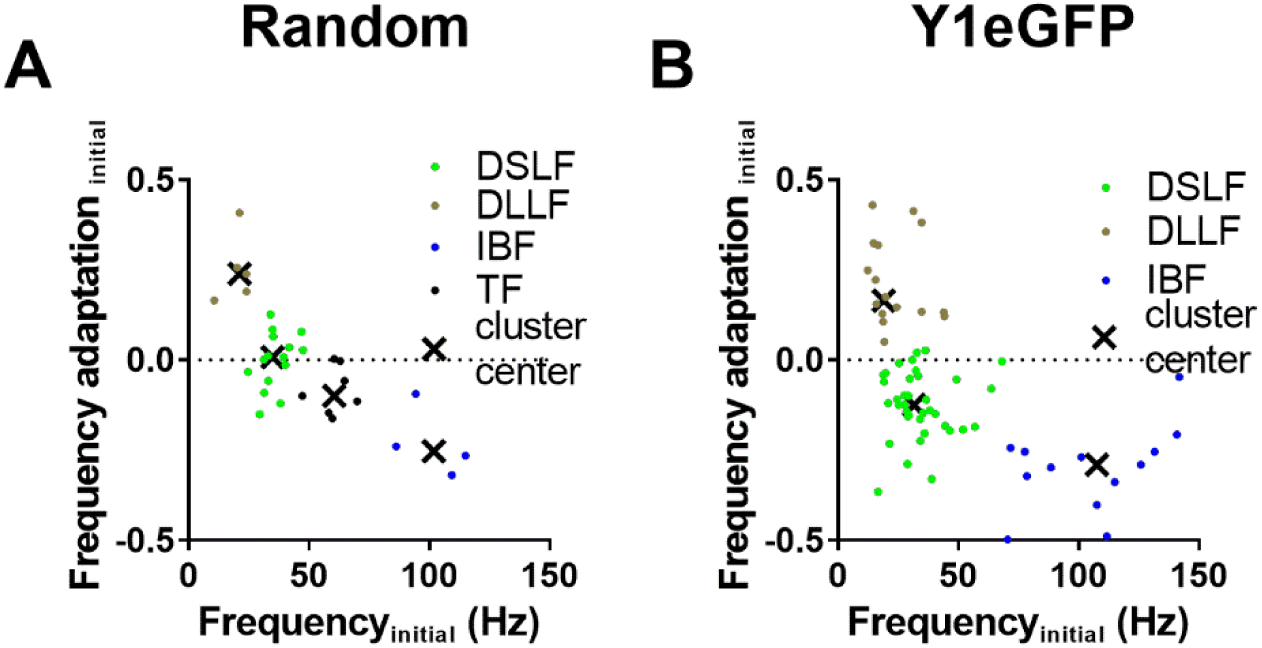
K-means cluster analysis-based grouping of firing patterns elicited from hyperpolarized conditions: K-means cluster analysis of (**A**) randomly sampled and (**B**)Y1eGFP neurons, in which two parameters – initial frequency and initial frequency adaptation – are used to assign each data point to one of the group based on how near they are to the respective mean. See associated Table 5 for prevalence of clustering. Each color represents a separate group and ‘X’ marks their mean center.

**Table 6:**
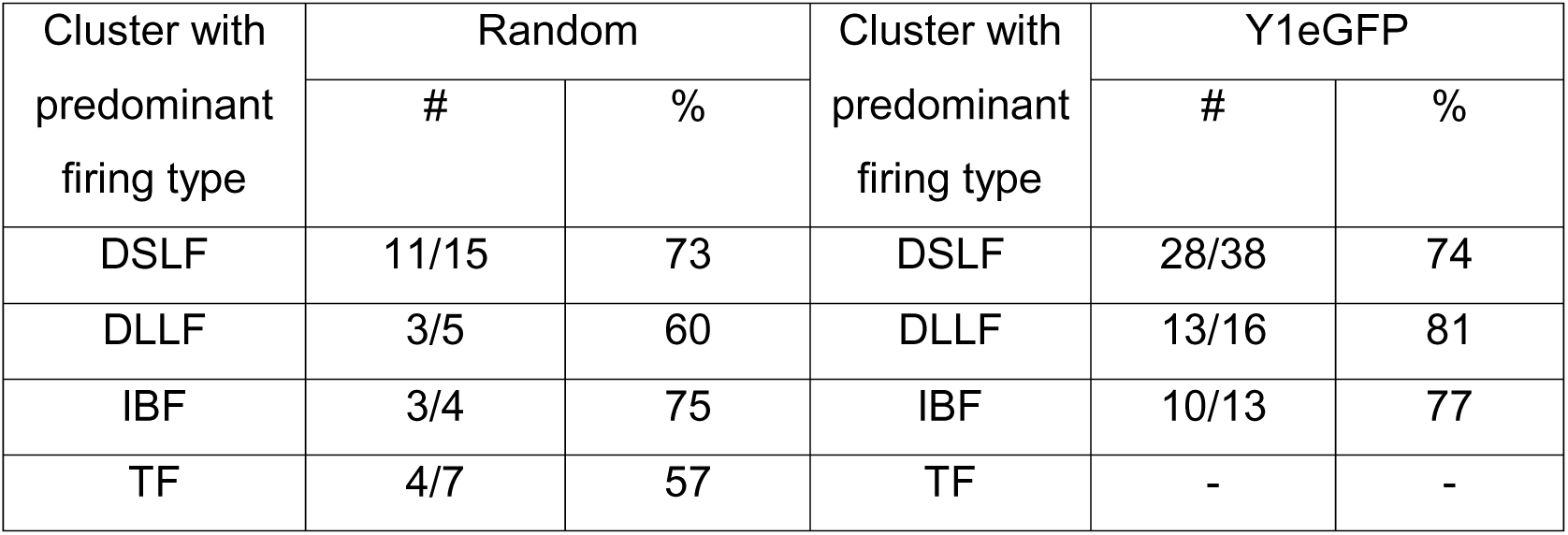
Neuronal populations in segregated clusters obtained by k-means cluster analysis for randomly sampled (Random) and Y1eGFP SDH neurons. Values indicate the number (#) or percentage (%) of neurons within each cluster. DSLF stand for delayed short latency firing, DLLF for delayed long latency firing, IBF for initial burst firing and TF for tonic firing.

### Rebound spiking occurred more frequently in Y1eGFP neurons as compared to randomly sampled neurons

Cessation of conditioning hyperpolarizing current can lead to rebound action potential firing (rebound spiking) in SDH neurons. Rebound spiking is likely mediated by T-type Ca channels (Candelas *et al*., 2019), and is typically observed as multiple action potentials in neurons with the IBF pattern (Cain & Snutch, 2013). Consistent with this literature, **Figure 9** illustrates rebound spiking in both randomly sampled and Y1eGFP neurons upon cessation of the 500 ms hyperpolarization current step. This varied with the subpopulation: **<u>DSLF</u>**, 6% (2 of 31) of randomly sampled, DSLF neurons (adapting and non-adapting) exhibited a single action potential on rebound from hyperpolarization (Figure 9A). By contrast, a much greater percentage of Y1eGFP DSLF neurons (32%, 18 of 56), exhibited a single action potential on rebound from hyperpolarization (P=0.007, χ test, Figure 9B); **<u>IBF</u>**, 16% (2 of 16) of randomly sampled, IBF neurons exhibited a burst of multiple action potentials on rebound from hyperpolarization (Figure 9C). A significantly greater percentage of Y1eGFP IBF neurons (48%, 10 of 21) exhibited burst of multiple action potentials on rebound from hyperpolarization (P=0.03, χ test, Figure 9D). At high current injection strengths, Y1eGFP IBF neurons exhibited fewer action potentials and higher firing frequency compared to randomly sampled IBF neurons (Table 3-5); **<u>PF</u>**, 57% (4 of 7) Y1eGFP neurons and no randomly sampled PF neurons exhibited single action potential on rebound from hyperpolarization; **<u>DLLF & TF</u>**, neither randomly sampled nor Y1eGFP neurons with DLLF or TF patterns exhibited rebound spiking.

**Figure 9.**
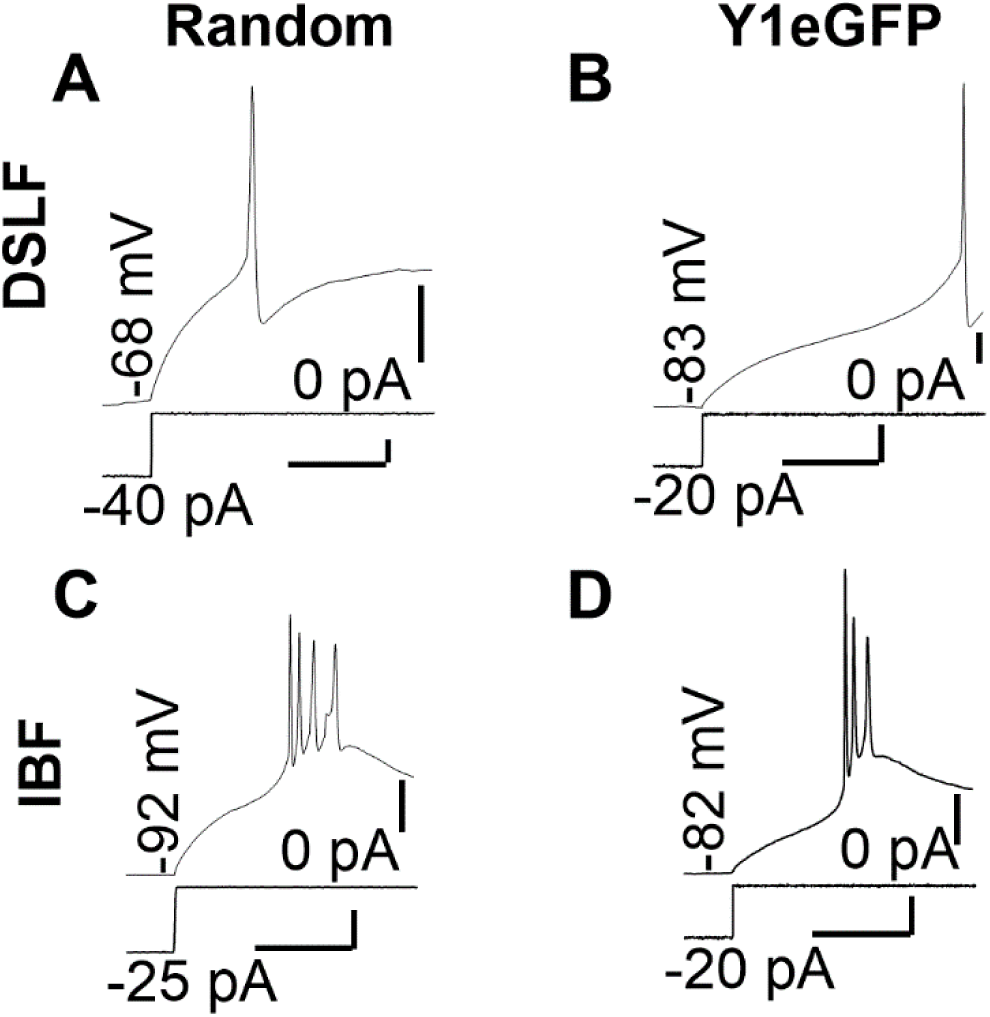
Two types of rebound spiking in randomly sampled (Random) and Y1eGFP neurons: **(A-D**) Action potentials firing observed after cessation of conditioning hyperpolarizing currents (with respective hyperpolarizing steps given below each graph) exhibited by (**A, C**) randomly sampled (Random) and (**B, D**) Y1eGFP neurons. 6% of randomly sampled (**A**) and 32% of Y1eGFP (**B**) DSLF (non-adapting and adapting) neurons exhibited single action potential on rebound from hyperpolarization. 16% of randomly sampled (**C**) and 48% of Y1eGFP (**D**) IBF neurons exhibited rebound spiking but with multiple action potentials. Scale bars: horizontal – 100 ms, vertical – 20 mV (above) and 15 pA (below).

### DSLF and DLLF neurons exhibited Fast and Slow A-type currents, respectively

#### Voltage activated transient outward currents

A-type K^+^ channels control the firing of action potential generation by delaying the onset of firing, which is observed as delayed firing in current clamp recordings (Hu *et al*., 2006; Hu & Gereau, 2011). Previous studies in spinal cord slices from adult rodents have segregated A-type currents into fast A-type currents (I_Af_) and slow A-type currents (I_As_) (Ruscheweyh *et al*., 2004; Yasaka *et al*., 2010; Smith *et al*., 2015). Although it seems possible that a faster decaying A-type current would be associated with a short-lived suppression (shorter latency or briefer delay) of the action potential firing during steady state current injection, it remained unclear whether I_Af_ and I_As_ are associated with the DSLF or DLLF phenotypes, respectively. To address this question, **Figure 10** used whole-cell patch-clamp recordings in voltage clamp mode to elicit voltage activated transient outward currents (TOC) in DSLF and DLLF neurons. **<u>DSLF</u>.** In randomly sampled and Y1eGFP DSLF neurons (both DSLF-N and DSLF-A), application of depolarizing voltage steps evoked TOCs in activating (Figure 10A-B) and inactivating steps (Figure 10C-D) below -40 mV (below threshold for activation of Na^+^ currents). The elicited TOC peaked within 9 milliseconds and subsequently decayed monotonically. The TOC decay constant for DSLF neurons was fast (∼ 40 ms for randomly sampled neurons and ∼ 30 ms for Y1eGFP neurons) for voltage steps from -80 mV to a sub-threshold range -55 to -40 mV (Table 7, left). The steady state (sustained) current amplitude after 250 ms of activation increased at a faster rate with increasing depolarization steps in randomly sampled than in Y1eGFP neurons (Table 7, left).

**Figure 10.**
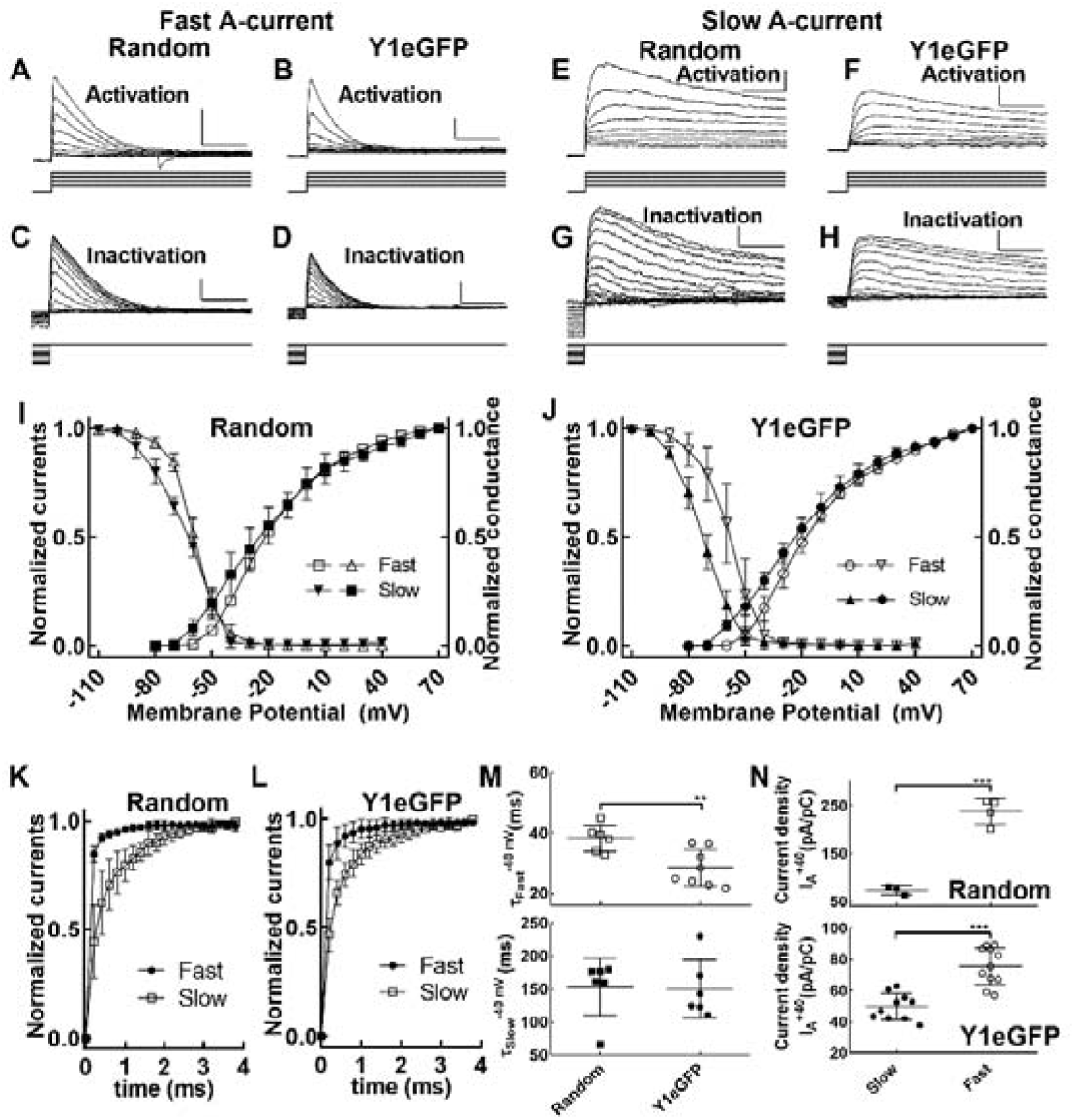
Voltage-dependent Fast A-currents in DSLF neurons and Slow A-type currents in DLLF neurons. Transient outward currents (TOCs) in response to a series of depolarizing voltage steps (-80 to -40 mV in +5 mV increments, shown at bottom) from -100 mV exhibited by (**A**) randomly sampled and (**B**) Y1eGFP neurons identified as Fast A-currents. In the same set of neurons, TOCs responses following a series of inactivating subthreshold depolarizing voltage steps (-110 and -40 mV in +5 mV increments, shown at bottom) to -40 mV in (**C**) randomly sampled and (**D**) Y1eGFP neurons. TOCs in response to activating steps (same steps as in A-B) exhibited by (**E**) randomly sampled and (**F**) Y1eGFP neurons identified as Slow A-currents. In the same set of neurons, TOCs in response inactivating steps (same steps as in **C-D**) in (**G**) randomly sampled and (**H**) Y1eGFP neurons. Scale bars: horizontal 200 ms and vertical 200 pA. (**I**) In randomly sampled neurons: activation curves (open squares) and inactivation curves (open triangles) obtained from isolated (in the presence of 2µM TTX, 10 mM TEA and 200 µM Cd^2+^) Fast A-type currents. In the same figure activation (closed squares) and inactivation (closed inverted triangles) curves obtained from isolated Slow A-type currents. (**J**) Similar curves obtained from TOCs in Y1eGFP neurons identified as Fast A-currents: activation (open circles), inactivation (open inverted triangles); and Slow A-currents: activation (closed circles), inactivation (closed triangles). (**K**) Recovery from inactivation for Fast (closed circles) and Slow (open squares) A-type currents in randomly sampled neurons. (**L**) Similar recordings from Y1eGFP neurons. (**M**) Decay constant obtained from Fast (top: RS, N=6 mice, 2 neurons were averaged in 2 of the mice; Y1eGFP: N=8 mice, 2 neurons were averaged for 3 of the mice; ** P=0.005: Student’s t-test, Random vs Y1eGFP) and Slow (bottom; RS and Y1eGFP: N=6 mice, 2 neurons were averaged in one mouse) A-type currents in randomly sampled and Y1eGFP neurons in response to steps from -100 to -40 mV. ** P<0.01: Student’s t-test, Random vs Y1eGFP. (**N**) Current density calculated for Fast (RS, N=4 mice, two neurons were averaged in one mouse; Y1eGFP, 11 mice, 2 neurons were averaged for one of the mice) and Slow (RS, 3 mice, one neuron per mouse; Y1eGFP, 11 mice, 2 neurons were averaged for one of the mice) A-type currents in randomly sampled (top) and Y1eGFP (bottom) neurons. *** P<0.001: Student’s t-test, Slow vs. Fast. Values in M-N represent mean ± SD; each dot represents average value of one mouse.

**Table 7:**
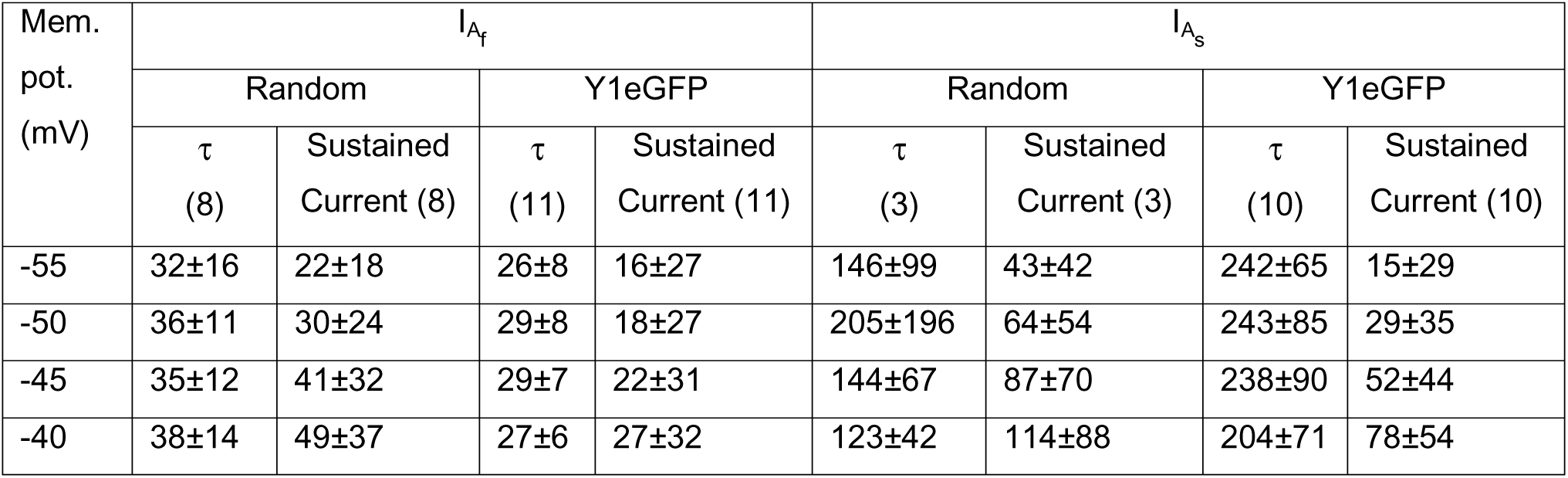
Duration (ms) of decay constant (τ) and sustained currents for randomly sampled (Random) and Y1eGFP superficial dorsal horn neurons exhibiting Fast (I_Af_) and Slow (I_As_) A-type currents in the absence of TEA, Cd2+ and TTX. Both decay constants and sustained currents were measured for currents elicited by voltage steps from -80 mV to a sub-threshold range between -55 to -40 mV (in +5 mV steps). τ was obtained by fitting a single exponent to the decay component of the transient outward current. Sustained currents were measured at 250 ms after activation (steady state). Values represent mean ± SD. Numbers in brackets indicates number of neurons from as many animals.

| Mem. pot. (mV) | $I_{A_f}$ | | | | $I_{A_s}$ | | | |
| --- | --- | --- | --- | --- | --- | --- | --- | --- |
|  | Random |  | Y1eGFP |  | Random |  | Y1eGFP |  |
| | $\tau$ (8) | Sustained Current (8) | $\tau$ (11) | Sustained Current (11) | $\tau$ (3) | Sustained Current (3) | $\tau$ (10) | Sustained Current (10) |
| -55 | 32±16 | 22±18 | 26±8 | 16±27 | 146±99 | 43±42 | 242±65 | 15±29 |
| -50 | 36±11 | 30±24 | 29±8 | 18±27 | 205±196 | 64±54 | 243±85 | 29±35 |
| -45 | 35±12 | 41±32 | 29±7 | 22±31 | 144±67 | 87±70 | 238±90 | 52±44 |
| -40 | 38±14 | 49±37 | 27±6 | 27±32 | 123±42 | 114±88 | 204±71 | 78±54 |

Further studies are needed to clarify this difference in the slow-rising, TEA sensitive K^+^ currents (I_K_), thought to contribute to sustained currents (Bourque, 1988). **<u>DLLF</u>**. The randomly sampled and Y1eGFP DLLF neurons exhibited voltage-activated TOCs in activating (Figure 10E-F) and inactivating steps (Figure 10G-H) with a slow decay constant. The elicited TOC decay constant for neurons was slow (> 100 ms for randomly sampled neurons and Y1eGFP neurons) for voltage steps from -80 mV to a sub-threshold range -55 to -40 mV (Table 7, right; Fig. 10M, below). The steady state (sustained) current amplitude after 250 ms of activation were larger in the DSLF randomly sampled than Y1eGFP neurons (Table 7, right).

#### Activation and inactivation curves

**<u>DSLF</u>**. In both randomly sampled (Figure 10I, open symbols) and Y1eGFP (Figure 10J, open symbols) DSLF neurons, the isolated fast TOCs I_Af_ exhibited activation (open squares) at about -50 mV and complete steady-state inactivation (open triangles) at -30 mV. Both randomly sampled (Figure 10K, filled symbols) and Y1eGFP (Figure 10L, filled symbols) neurons exhibited a fast recovery after inactivation that was back to normal within a few msec. **<u>DLLF</u>**. In randomly sampled (Figure 10I, filled symbols) and Y1eGFP (Figure 10J, filled symbols) DLLF neurons, the isolated slow TOCs I_As_, exhibited activation at a more hyperpolarizing potential (∼-60 mV) compared to I_Af_ and inactivation at ∼-40 mV for randomly sampled (filled inverted triangles) and at approximately -30 mV for Y1eGFP neurons (filled triangles). The randomly sampled (Figure 10K, open symbols) and Y1eGFP (Figure 10L, open symbols) neurons exhibited a slow recovery after inactivation that lasted up to approximately 3 sec.

In randomly sampled neurons, the slope of the slow activation curves was significantly shallower compared to the fast activation curves (p = 0.04; Table 8; Statistical Summary Table, P6). Among the Y1eGFP neurons, V_1/2_ for I_As_ for fast activation curves was significantly depolarized (p=0.01; Table 8) and exhibited a significant steeper slope (p=0.02; Table 8) compared to the V_1/2_ for I_Af_ for slow activation curve. Similarly, V_1/2_ for I_As_ for fast inactivation curves was significantly depolarized (p =0.0001; Table 8) and exhibited a significantly steeper slope (p= 0.04; Table 8) compared to the slow I_Af_ inactivation curve.

**Table 8:** Half-maximal voltage (*V_1/2_*) and slope factor (*k*) obtained by fitting fast and slow activation and inactivation curves of A-type currents to Boltzmann function for randomly sampled (Random) and Y1eGFP SDH neurons. Numbers in brackets indicates number of neurons from as many animals. Values represent mean ± SD. Groups are indicated within square brackets: a) *V_1/2_* and b) *k* for randomly sampled fast activation/inactivation curves; c) *V_1/2_* and d) *k* for randomly sampled slow activation/inactivation curves; e) *V_1/2_* and f) *k* for Y1eGFP fast activation/inactivation curves; g) *V_1/2_* and h) *k* for Y1eGFP slow activation/inactivation curves. *p<0.05, **p<0.01 or ***p<0.001: Student’s t-test between groups shown in curly brackets.

|  | Random |  |  |  | Y1eGFP |  |  |  |
| --- | --- | --- | --- | --- | --- | --- | --- | --- |
|  | Fast (n=6) |  | Slow (n=3) |  | Fast (n=9) |  | Slow (n=6) |  |
| | [a] half maximal voltage ($V_{1/2}$) | [b] Slope (k) | [c] half maximal voltage ($V_{1/2}$) | [d] Slope (k) | [e] half maximal voltage ($V_{1/2}$) | [f] Slope (k) | [g] half maximal voltage ($V_{1/2}$) | [h] Slope (k) |
| Activation curve | -18.3±1.8 | 16.7±0.7 | -17.0±2.7<br>{a,c; ns. } | 21.1±3.4<br>{b,d; *} } | -15.0±3.2<br>{a,e; *} } | 17.9±1.5<br>{b,f; ns. } | -20.4±3.9<br>{c,g; ns. }<br>{e,g; *} } | 20.5±2.0<br>{d,h; ns. }<br>{f,h; *} } |
| Inactivation curve | -59.4±1.6 | 6.6±1.2 | -63.9±1.7<br>{a,c; *} } | 10.0±1.7<br>{b,d; *} } | -58.5±6.1<br>{a,e; ns. } | 7.0±0.9<br>{b,f; ns. } | -72.5±2.6<br>{c,g; **}<br>{e,g; ***} | 8.0±0.8<br>{d,h; ns. }<br>{f,h; *} } |

#### Y1eGFP DSLF-A neurons exhibit faster A-type currents with shorter latency and smaller current density as compared to randomly sampled DSLF-N neurons

Figures 5-6 indicate that Y1eGFP neurons are enriched in DSLF-A. Furthermore, the latency of action potential firing to onset of current injection (delay) was shorter in Y1eGFP DSLF-A neurons compared to randomly sampled DSLF-N neurons (Table 3). Because A-currents are thought to be responsible for the DSLF phenotype (Hu *et al*., 2006), we hypothesized that the decay constant of A-currents would be smaller in Y1eGFP neurons. Indeed, a TOC with faster decay kinetics that underlies DSLF firing exhibited characteristically shorter latency to first spike for Y1eGFP DSLF-A neurons compared to randomly sampled DSLF-N neurons (Table 3 and Fig. 10M above, p<0.005, Statistical Summary Table, P7). The characteristics of I_As_ of randomly sampled neurons were, however, similar to Y1eGFP DLLF neurons (Table 3 and Figure 10M below, p=0.9, Statistical Summary Table, P7).

The current densities for isolated I_Af_ and I_AS_ were larger in randomly sampled neurons as compared to Y1eGFP neurons, (Figure 10N: compare respective values for I_Af_ and I_AS_ between top and bottom sub-figures, P<0.00001 (I_Af_), P = 0.001 (I_AS_), Statistical Summary Table, P8) suggesting a higher expression of channels mediating A-currents in randomly sampled neurons as compared to Y1eGFP neurons.

#### Pharmacological segregation of Fast and Slow A-currents using the K_v_4 channel blocker, 4-AP

**Figure 11** illustrates the sensitivity of TOCs to 4-aminopyridine (4-AP), which is widely used to inhibit the K_v_4 channels that mediate I_Af_ and I_As_ (Ruscheweyh & Sandkühler, 2002; Ruscheweyh *et al*., 2004; Yasaka *et al*., 2010; Smith *et al*., 2015). The lower concentration of 4-AP (0.5 mM) partially reduced slow A-currents while sparing fast A-currents; by contrast, the higher concentration (5 mM) blocked both slow and fast A-currents, allowing us to pharmacologically segregate them (Ruscheweyh *et al*., 2004). **<u>Fast A-currents in DSLF neurons:</u>** In the presence of TTX, TEA and Cd^2+^, the effect of 0.5 mM 4-AP on I_Af_ was minimal, but 5 mM 4-AP abolished fast TOCs (Figure 11A). Y1eGFP DSLF-A neurons responded to 4-AP in a fashion similar to randomly sampled neurons (Figure 11B). **<u>Slow A-currents in DLLF neurons:</u>** In the presence of TTX, TEA and Cd^2+^, 4-AP acted on I_As_ in a concentration-related manner in randomly sampled (Figure 11C) and Y1eGFP (Figure 11D) neurons.

**Figure 11.**
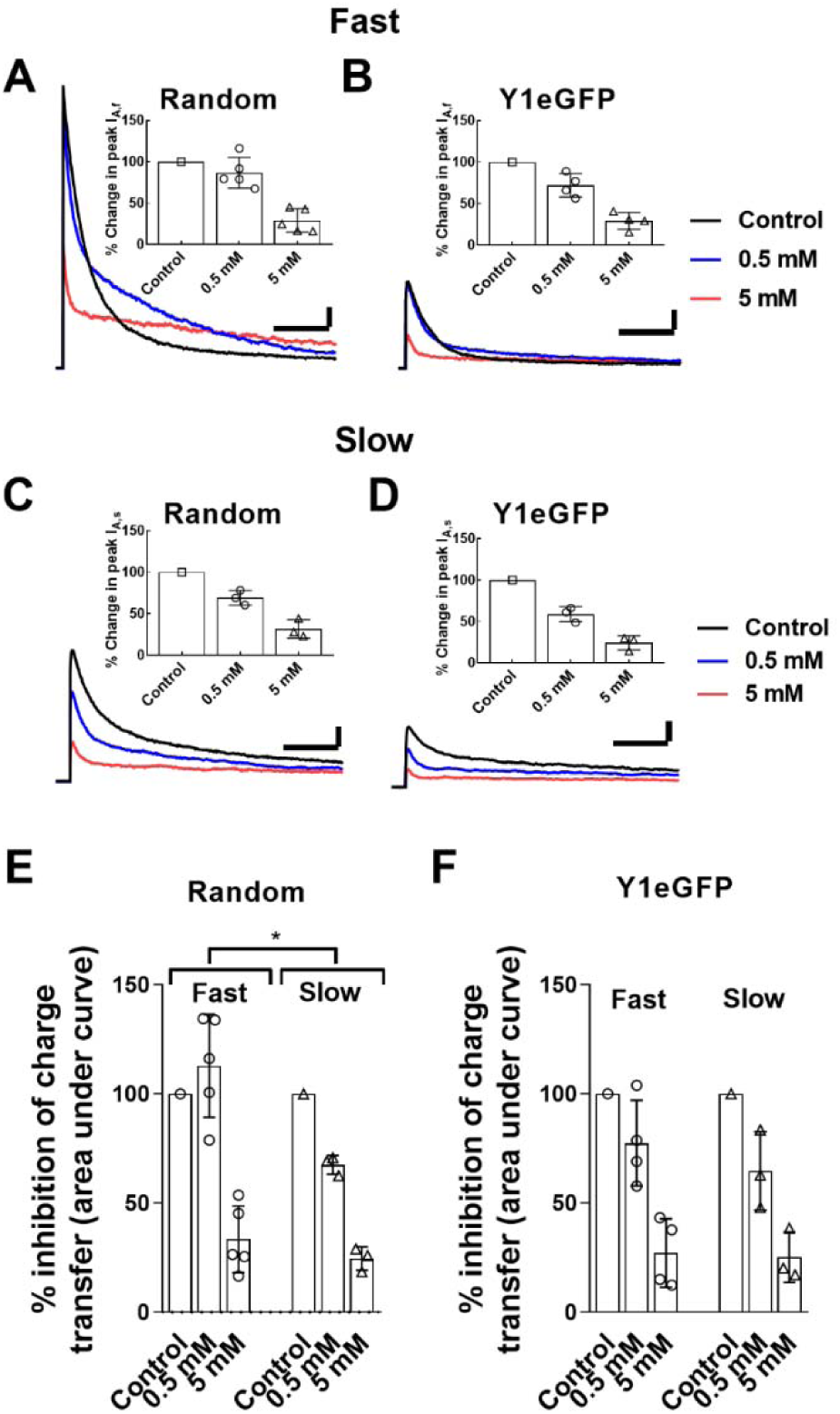
Inhibition of the Fast and Slow A-type currents by 4-AP in SDH neurons. Examples of inhibition by 4-AP of the isolated Fast A-type currents exhibited by (**A**) randomly sampled and (**B**) Y1eGFP neurons in the presence of 2µM TTX, 10 mM TEA and 200 µM Cd^2+^. Insets: summary of their % change in peak currents induced by 4-AP (RS, N=5 mice, 8 neurons; Y1eGFP, N=4 mice, 10 neurons). Similar Inhibition of the isolated Slow A-type currents exhibited by randomly sampled (**C**, inset) and Y1eGFP (**D**, inset) neurons (RS and Y1eGFP: N=3 mice, one neuron per mouse). Scale bars: horizontal – 100 ms, vertical – 500 pA. Summary of inhibition by 0.5 mM and 5 mM 4-AP to Fast and Slow A-type currents, expressed as percentage change in area under the curve in randomly sampled (**E**) and Y1eGFP (**F**) neurons. Values in A – D (insets), E and F represent mean ± SD; each dot represents average value per one mouse. *p=0.02: 2-way ANOVA, 4-AP conc. vs A-current type.

We observed a statistically significant relationship between the type of A-currents (slow and fast) and drug concentration, finding an interaction for area under the curve between the type of A-current and drug concentration (P=0.02, 2-way ANOVA; Statistical Summary Table, P9) in the randomly selected neuron group (Figure 11E). These data suggest that 4-AP inhibits the fast and slow A-currents differently. Thus, the TOCs exhibited by delayed firing, randomly sampled neurons express A-type potassium currents with slower (*I_As_*) or faster (I_Af_) kinetics, correlated with 4-AP sensitivity. This suggests the TOCs are mediated by K_v_4 channels comprised of different subunits. There was no statistical significance between the type of A-currents (slow and fast) and drug concentration (P>0.05, 2-way ANOVA) for Y1eGFP cells (Figure 11F), suggesting less variability in K_v_4 channel subunit composition in these neurons.

### T-type calcium currents underlie Initial burst firing patterns

In most cases, randomly sampled and Y1eGFP neurons with IBF firing exhibited an inward current in response to steps from -100 mV to -50 mV (Figure 12A-B), whereas a TOC would typically be elicited in l delayed firing neurons. **Figure 12** illustrates that bath application of 100 µM Ni^2+^ abolished the inward currents in both IBF randomly sampled and Y1eGFP neurons (respective insets in Figure 12A-B). These inward currents evoked in response to voltage steps from -100 mV to -50 mV are mediated by T-type Ca^2+^ channels and are referred to as *I_Ca,T_*, consistent with previous reports (Ruscheweyh *et al*., 2004; Yasaka *et al*., 2010; Smith *et al*., 2015).

**Figure 12.**
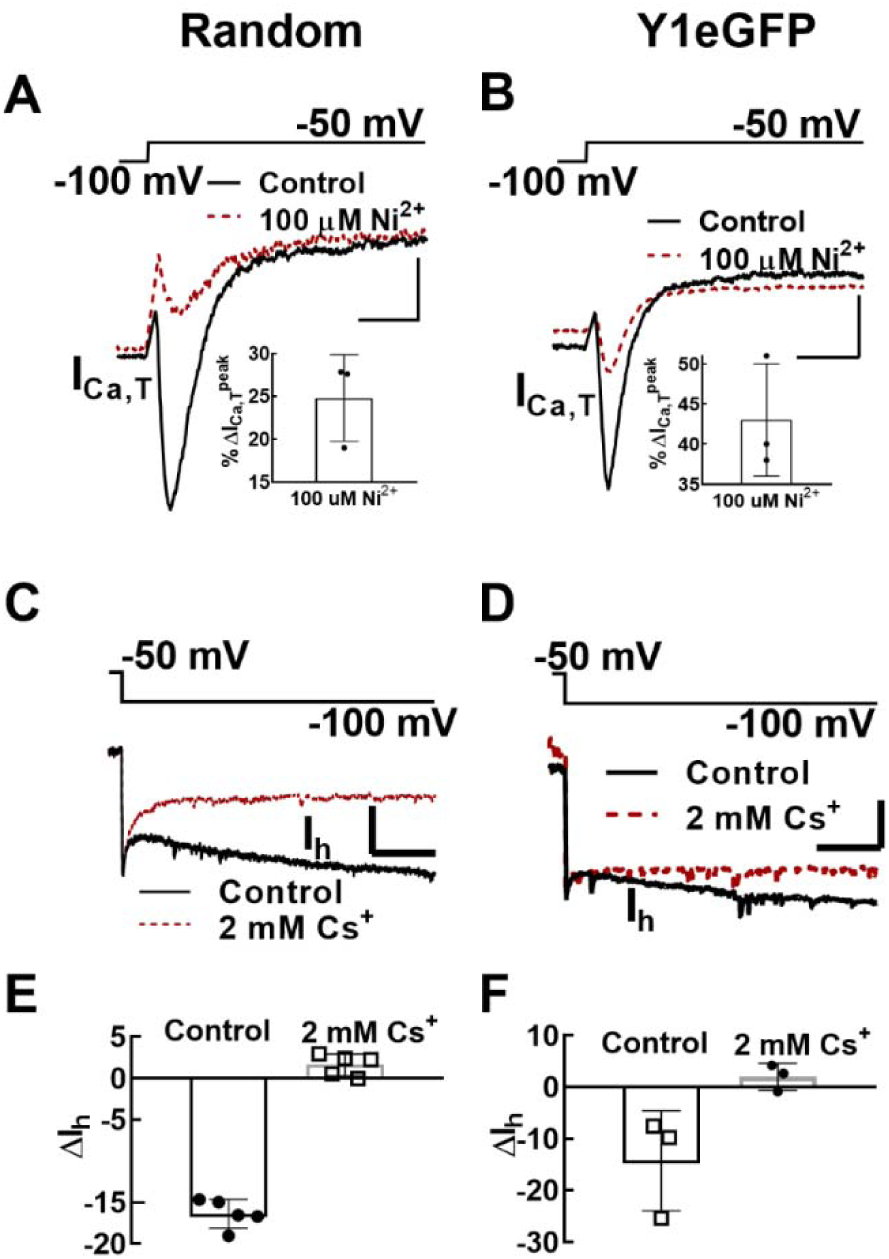
Voltage activated currents exhibited by initial burst firing (IBF) and tonic firing (TF) neurons. (**A**) Step-up potential from -100 mV to low thresholds (-50 mV) induced inwardly directed currents (*I_Ca,T_*) in randomly sampled IBF neurons. Inset: Quantification of the relative block by Ni^+^ for respective samples (N=3 mice, one neuron per mouse). (**B**) Similar behavior is observed in Y1eGFP IBF neurons (inset: relative block by Ni^+^; N=3 mice, one neuron per mouse). Scale bars: horizontal – 40 ms, vertical – 40 pA. (**C**) Hyperpolarizing step-down potential from -50 mV to -100 mV activated slow inward current (*I_h_*) in randomly sampled tonic firing neurons (**D**) Similar behavior is observed in Y1eGFP neurons. Scale bars: horizontal – 200 ms, vertical – 20 pA. (**E**) Quantification of the relative (completely) block by 2 mM Cs^+^ of *I_h_* in randomly sampled neurons (N=5 mice, 6 neurons). (**F**) Similar relative block in Y1eGFP neurons (N=3 mice, one neuron per mouse). Values in A (inset), B (inset), E and F represent Mean ± SD; each dot represents average value of one mouse.

### Hyperpolarization-activated inward currents and Tonic firing

Some dorsal horn interneurons express hyperpolarization-activated currents (*I_h_*) mediated by a family of non-selective cation channels (i.e., HCN channels) (Gold & Gebhart, 2010; Yasaka *et al*., 2010; Hughes *et al*., 2012, 2013; Smith *et al*., 2015). Consistent with these previous reports, in 50% of TF randomly sampled and Y1eGFP neurons, a slow hyperpolarization activated inward current was observed during 1 s steps from -50 mV to -100 mV (Figure 12C-D). These slow inward currents were sensitive to 2 mM Cs^+^ in both randomly sampled and Y1eGFP neurons (Figure 12E-F). TF neurons, mostly in randomly sampled neurons, also expressed the most robust *I_h_* currents. *I_h_* currents are often associated with SDH “islet cells” that have an extensive dendritic spread (Yasaka *et al*., 2010; Smith *et al*., 2015). However, a few of the IBF neurons also exhibited both *I_Ca,T_* and *I_h_*.

## DISCUSSION

Immunohistochemical, in situ hybridization, and single-cell RNA sequencing studies indicate that the vast majority of Y1R-expressing interneurons (Y1-INs) are glutamatergic (Nelson & Taylor, 2020). For example, Y1-INs express markers of excitatory interneurons in the dorsal horn, including somatostatin and Tlx3 (Zhang *et al*., 1999; Häring *et al*., 2018; Sathyamurthy *et al*., 2018; Dickie *et al*., 2019; Nelson *et al*., 2019; Nelson & Taylor, 2020). Furthermore, neuronal ablation studies indicate that Y1-INs contribute to the behavioral and molecular signs of inflammatory and neuropathic pain (Wiley *et al*., 2009; Nelson *et al*., 2019). Our *in situ* hybridization experiments in Y1eGFP reporter mice confirmed Y1 mRNA expression in the great majority of GFP positive SDH neurons. With this powerful technique, the current studies reveal multiple subpopulations of Y1-INs based on their electrophysiological properties - information that will become quite useful as we learn more about their contribution to nociception and chronic pain.

### Y1R-selective agonists induce outward currents, hyperpolarize the membrane potential, and inhibit stimulus-evoked action potentials in Y1eGFP neurons

Anatomical and pharmacological data led to the speculation that NPY, released from inhibitory interneurons, can modulate spinal pain modulatory neurons in SDH that express the NPY Y1 receptor (Zhang *et al*., 1999; Brumovsky *et al*., 2006; Smith *et al*., 2007). The first neurophysiological tests of this hypothesis came from early studies in unidentified dorsal horn interneurons in rat spinal cord slices (Moran *et al*., 2004; Miyakawa *et al*., 2005; Melnick, 2012). For example, Miyakawa *et al* (2005) reported that NPY elicited outward currents, but because these studies were limited to unidentified neurons under voltage clamp conditions, their relevance to the prevailing conceptual mechanisms of NPY-mediated pain modulation remained unclear (Miyakawa *et al*., 2005). To address this question, we recorded under both voltage-clamp and current-clamp conditions in Y1-eGFP neurons. Furthermore, we extended previous studies with evaluation of the effects not only of NPY but also of a Y1R-selective agonist, and we evaluated their ability to modulate primary afferent stimulus-evoked action potentials. Either NPY or the Y1R-selective agonist [Leu^31^,Pro^34^]-NPY elicited outward currents and membrane hyperpolarization.

An important caveat here is that, although GFP expression coincided with Y1 mRNA transcription indicating high fidelity in Y1 receptor-expressing neurons, hyperpolarization was observed in less than half of all Y1eGFP neurons tested. Notably, none of the IB firing neurons were responsiveness to NPY. We cannot exclude the possibility that insertion of eGFP might reduce functional responsiveness of the Y1 receptor to NPY. The transgene could be expressed transiently during early development or cause developmental changes or steric hindrance, leading to interruption of receptor trafficking (including receptor internalization), receptor availability at the plasma membrane, or number of functional receptors.

Regardless, the available responses of Y1eGFP neurons to Y1R agonists indicate that they are largely restricted to outward currents and hyperpolarization. Together with our finding that Y1R agonist suppressed dorsal root stimulus-evoked firing of action potentials in Y1eGFP neurons, we conclude that NPY inhibits neuronal excitability, including in Y1R expressing neurons that receive primary sensory afferent input.

### NPY produces inwardly rectifying K^+^ currents in Y1eGFP neurons

Inwardly-rectifying K^+^ channels (K_ir_), also known as G protein-coupled inwardly-rectifying K^+^ (GIRK) channels, contribute to membrane inhibition and thus control the excitability of the neuron. GIRK channels that likely mediate the ability of Y1R agonists to elicit inward rectification have been measured previously using a voltage step protocol on unidentified rat SDH neurons (Moran *et al*., 2004; Miyakawa *et al*., 2005). Consistent with these results, we found that NPY and a Y1R agonist elicited comparable inward rectification in identified mouse Y1eGFP neurons. Furthermore, to obtain a more convincing estimate of inward rectification, we then went beyond previous studies with evaluation of conductance and quantification of the degree of inward rectification. We found that K^+^ ions pass more easily (2-fold) into the cell as compared to out of the cell. Our values are also consistent with the degree of inward rectification of NPY- sensitive currents seen in other systems such as C-cells of amphibian sympathetic ganglia, rat arcuate nucleus, and rat thalamic neurons (Zidichouski *et al*., 1990; Sun & Miller, 1999; Sun *et al*., 2001). Moran et. al., 2004, reported that NPY acted on Y1R on presynaptic terminals of interneurons in SDH to suppress GABA release and reduce inhibitory currents (Moran *et al*., 2004). Our observation of NPY induced inwardly rectifying currents suggests an additional, postsynaptic mechanism that might compete against the Y1R-mediated attenuation of inhibitory synaptic transmission. In summary, the neurophysiological actions of Y1R-selective agonists on Y1eGFP neurons of the SDH (outward currents, hyperpolarization of the membrane potential, inhibition of stimulus-evoked action potentials, inward rectification through GIRK channels) are consistent with NPY mediated inhibition.

#### NPY targets K_v_4 channels on Y1R-expressing interneurons to inhibit nociception

K_v_4.2 mediates the A-type current in SDH neurons and can regulate the neuronal excitability and central sensitization that underlies hypersensitivity after tissue injury (Hu *et al*., 2006). The NPY mediated hyperpolarization of the pronociceptive Y1-expressing neurons (Nelson *et al*., 2019) would be expected to shift PF and TF neurons to delayed firing (Fig. 6), possibly by unmasking of the A-type currents. In addition to reducing responsiveness to afferent stimulation (Fig. 3I), engaging K_v_4 that mediates delayed firing could make them less pronociceptive by limiting action potential firing. Thus we suggest that nerve injury-induced upregulation of NPY reduces the excitability of excitatory Y1R-expressing neurons by both hyperpolarizing the membrane and by unmasking TOCs, and this decrease in excitability contributes to NPY mediated anti-nociception (Brumovsky *et al*., 2007; Todd, 2017; Nelson *et al*., 2019).

### Y1eGFP neurons exhibit phasic rather than delayed firing when studied at or near resting membrane potential

Literature from the last fourteen years describes the electrophysiological properties of genetically-defined subpopulations of interneurons in the dorsal horn (Schoffnegger *et al*., 2006; Punnakkal *et al*., 2014; Smith *et al*., 2015), and we extended this database to Y1R-expressing neurons. Furthermore, a combined current-clamp electrophysiological and anatomical approach is increasingly used to segregate subpopulations of inhibitory and excitatory interneurons in SDH (Yasaka *et al*., 2010; Punnakkal *et al*., 2014; Smith *et al*., 2015), and in this vein we sought to define the firing patterns of Y1-INs. Our profiling of Y1eGFP neurons found some similarities to randomly sampled SDH neurons such as membrane capacitance, membrane resistance and resting membrane potential. On the other hand, Y1eGFP neurons exhibited a less negative action potential threshold, indicative of a potential for decreased excitability in response to, for example, excitatory synaptic input. They also exhibited a longer action potential width at base, which would tend to reduce action potential rate, but could also reflect increased calcium influx, which could have multiple effects on cellular excitability (Bean, 2007).

When studied at or near resting membrane potential, randomly sampled neurons frequently exhibited five major firing patterns, with the majority being delayed firing (DSLF 46%, DLLF 13%). Y1eGFP neurons were mostly PF, being much less likely to exhibit delayed firing (12%). This lack of delayed firing in excitatory Y1eGFP neurons contrasts with the preponderance of delayed firing observed in other subpopulations of excitatory neurons in the SDH, including calretinin-expressing neurons (Smith *et al*., 2015), vGlut2-eGFP neurons (Punnakkal *et al*., 2014), and substance P-expressing neurons (Dickie *et al*., 2019). Our results are not the first, however, to counter the hypothesis that delayed firing is an essential characteristic of excitatory interneurons in the SDH (when studied at near resting membrane potential). For example, delayed firing has been observed in one-third of GAD67-eGFP inhibitory interneurons (Heinke *et al*., 2004; Schoffnegger *et al*., 2006; Hu & Gereau, 2011). Instead of delayed firing, we found that Y1eGFP neurons in the SDH were enriched in the PF phenotype. We conclude that Y1R-expressing interneurons represent a subclass of excitatory interneurons that predominantly exhibit PF, rather than delayed firing, when evaluated at resting membrane potential.

High-throughput, unbiased, transcriptomic analyses have revolutionized the segregation of interneuron populations in the dorsal horn (Häring *et al*., 2018; Sathyamurthy *et al*., 2018; Gatto *et al*., 2019; Nelson & Taylor, 2020; Peirs *et al*., 2020), promoting fluorescence *in situ* hybridization and immunohistochemical identification of genetic markers of excitatory interneurons (Dickie *et al*., 2019; Gutierrez-Mecinas *et al*., 2019; Bell *et al*., 2020; Peirs *et al*., 2020), including *Npy1r* as noted in our recent review (Nelson & Taylor, 2020). Notably, excitatory *Npy1r*-expressing neurons define the DE-2 (Sathyamurthy *et al*., 2018) and Glut8 clusters (Häring *et al*., 2018), both of which are also defined by the expression of *Grp* (codes for gastrin releasing peptide). Indeed, we recently used double fluorescence in situ hybridization to demonstrate that Y1eGFP interneurons colocalize with *Grp* mRNA (Nelson & Taylor, 2020). Interestingly, GRPeGFP neurons share neurophysiological properties with our Y1eGFP neurons in the rodent spinal cord slice (Koga *et al*., 2011; Dickie *et al*., 2019). For example, the DSLF and PF firing patterns of Y1eGFP neurons are similar to the phasic-like firing pattern that is exhibited by 50% of all GRPeGFP neurons when studied at resting membrane potential (Dickie *et al*., 2019). Furthermore, the proportion of GRPeGFP and Y1eGFP cells that exhibit subthreshold voltage gated currents are similar. The percentage of cells that exhibit I_Af_, I_As_, I_CaT_ and I_h_ currents: GRPeGFP = 41, 26, 33, and 37; Y1eGFP = 51, 26, 22 and 23, respectively.

### A preconditioning membrane hyperpolarization dramatically shifts Y1eGFP neurons to delayed firing phenotypes

To determine firing patterns in SDH neurons, current clamp recordings are typically initiated from a resting membrane potential that is above (less negative) than -75 mV (Ruscheweyh & Sandkühler, 2002; Ruscheweyh *et al*., 2004; Heinke *et al*., 2004; Balasubramanyan *et al*., 2006; Yasaka *et al*., 2010; Punnakkal *et al*., 2014; Smith *et al*., 2015; Stebbing *et al*., 2016). However, Schoffnegger and colleagues reported that gap firing, a type of delayed firing, is readily observed when current is applied when SDH neurons are held for at least 500 ms at potentials more negative than -75 mV (Schoffnegger *et al*., 2006). To test the hypothesis that we could capture delayed firing in Y1-eGFP neurons, we studied firing patterns that could be evoked not only at conditioning potentials that included depolarized potentials between -50 and -65 mV and near resting potential between -65 and -80 mV, but also at hyperpolarized potentials between -80 to -95 mV (Yasaka *et al*., 2010). When studied at these conditioning hyperpolarized potentials, we found that randomly sampled neurons frequently exhibited DSLF, DLLF, TF, and IBF firing patterns, and only rarely exhibited PF patterns (Figure 7). Y1eGFP neurons frequently exhibited DSLF, DLLF, and IBF, but only rarely exhibited TF or PF. Thus, our current clamp recordings in Y1eGFP neurons revealed a dramatic shift away from PF (from 59% at resting membrane potential conditions to 6% at hyperpolarized conditions) and towards an enrichment of delayed firing (from 12% delayed firing at resting membrane conditions to 70% delayed firing at hyperpolarized conditions). Remarkably, when comparing Y1eGFP neurons tested at resting membrane versus hyperpolarized potentials, we discovered that the PF and TF patterns were essentially replaced with the DSLF and DLLF patterns, respectively (Figure 6). A-type currents are voltage-gated potassium (K_v_) currents that activate after hyperpolarizing the cell membrane (V_m_ < -80 mV). When action potentials are evoked from hyperpolarized conditions, A-currents, expressed by three-fourth of Y1eGFP neurons, are activated and their expression manifests as delayed firing. Thus, analysis of action potential firing patterns evoked from hyperpolarized conditions revealed a dramatic occurrence of the DSLF firing pattern that would otherwise have been completely missed and could have been alternatively classified as phasic firing.

### Y1eGFP but not randomly sampled DSLF neurons are rapidly adapting

Our in-depth quantitative analysis of firing patterns revealed several quantitative differences between Y1eGFP as compared to randomly sampled neurons, including shorter 1^st^ spike latency, lower spike frequency, and fewer total number of action potentials. But most strikingly, a subset of Y1eGFP DSLF neurons exhibited spike-frequency adaptation (Benda & Herz, 2003) reminiscent of the spike-frequency adapting SDH neurons described in earlier studies (Grudt & Perl, 2002; Ruscheweyh & Sandkühler, 2002; Hu *et al*., 2003), some of which may represent nociceptive-specific SDH neurons (Lopez-Garcia & King, 1994). We introduce the acronyms DSLF-A and DSLF-N, respectively, to distinguish rapidly adapting neurons from the more typical non-rapidly adapting phenotype that are frequently observed in randomly sampled SDH neurons. Quantification of firing pattern frequency revealed that delayed firing Y1eGFP neurons were 47% DSLF-A and only 3% DSLF-N, while randomly sampled neurons were only 7% DSLF-A and 47% DSLF-N. Further characterization of Y1eGFP DSLF-A neurons revealed that they exhibit: (1) fast A-currents that decay faster than *I_Af_* in randomly sampled neurons; (2) smaller *I_Af_* density than randomly sampled neurons; (3) shorter delay compared to other firing type neurons; and (4) a higher percentage of rebound spiking, as discussed next. These features of Y1 expressing neurons, especially DSLF neurons, may be important determinants of nociceptive processing. In the setting of injury, for example, subtle differences in currents responsible for spike frequency adaptation and in the kinetics of the A-type current may substantially contribute to neuronal excitability and thus hypersensitivity.

### Rebound spiking is prevalent in the DSLF subpopulation of Y1eGFP neurons

Compared to randomly sampled neurons, the present studies indicate that a high percentage of Y1eGFP DSLF-A interneurons in the SDH exhibit rebound spiking. These rebound action potentials are likely mediated by T-type Ca^2+^ channels, as suggested by recent studies showing that the T-type channel inhibitor TTA-A2 greatly reduced the amplitude of rebound spiking in adult mouse SDH neurons (Candelas *et al*., 2019). Cav3.2 T-type calcium channels shift the intrinsic membrane properties of SDH neurons towards increased excitability (Candelas *et al*., 2019). Cav3.2 T-type calcium channels are highly expressed in SDH interneurons (60%) and likely participate in their increased excitability during conditions of central sensitization such as peripheral nerve injury (Feng *et al*., 2019). Furthermore, selective disruption of Cav3.2 channels in primary afferent nociceptors reduced behavioral signs of post-surgical pain (Joksimovic *et al*., 2018). We can speculate that outward, fast A-currents (I_Af_), which we found to be enriched in Y1eGFP DSLF-A neurons, operate in opposition with inward Cav3.2 currents (I_CaT_.) during action potential firing conditions. If so, then interventions that down-regulate A-type currents, such as nerve injury (Hu *et al*., 2006; Anderson *et al*., 2010) may unmask the ability of I_CaT_ to favor a state of increased excitability of DSLF-A Y1eGFP SDH interneurons. Because Y1R-expressing neurons contribute to neuropathic pain (Nelson *et al*., 2019), we suggest that the Y1eGFP DSLF-A sub-population is a potential target for the pain inhibitory actions of Cav3.2 T-type calcium channel blockers (Todorovic & Jevtovic-Todorovic, 2011).

### Voltage-dependent segregation of delayed firing neurons into DSLF with fast A-currents and DLLF with slow A-currents

The firing patterns assessed in current clamp mode are shaped by ionic conductance (Baxter & Byrne, 1991; Bean, 2007). An important example is the contribution of A-type K^+^ currents to delayed firing (Russier *et al*., 2003; Hu *et al*., 2006); A-currents control the rate of action potential generation by delaying the onset of firing (Hu & Gereau, 2011). Consistent with previous studies in superficial dorsal horn (Grudt & Perl, 2002; Yasaka *et al*., 2010; Smith *et al*., 2015), we identified four of the major types of voltage-activated currents in SDH neurons: 1) a 4-AP sensitive voltage activated fast outward A-current (I_Af_) (∼40 ms decay constant; 2) a 4-AP sensitive voltage activated slow outward A-currents (I_As_) (>100 ms decay constant); 3) a Ni^2+^ sensitive low threshold voltage gated inward T-type calcium current (I_Ca,T_); and 4) a Cs^+^ sensitive, hyperpolarization-activated slow inward current (I_h_). Of the multiple Kv families (K_v_1.4, K_v_3.4, K_v_4.1, K_v_4.2, and K_v_4.3) that can contribute to A-type currents in mammals (Coetzee *et al*., 1999; Carrasquillo & Nerbonne, 2014), K_v_4.2, and K_v_4.3 are known to be expressed in “pain excitatory” interneurons (Huang *et al*., 2005; Hu *et al*., 2006; Hu & Gereau, 2011; Kuo *et al*., 2017). Our results are consistent with multiple A-type currents due to the differences in decay time constant and 4-AP sensitivity. The fast and slow A-currents could be separated because: the membrane potential at which they activate, (ii) their characteristics of the activation and inactivation curve, (iii) the duration of recovery of their currents after inactivation, and (iv) their current densities were different. More importantly, we discovered that each type of voltage-activated current was closely associated with a specific firing pattern: DSLF exhibit I_Af_; DLLF exhibit I_As_; most IBF neurons exhibit I_Ca,T_ and half of TF neurons exhibit I_h_ currents. The correlation between firing pattern and ionic conductance (I_Af_, I_As_, I_Ca,T_ and I_h_) increases our understanding of how the expression of particular channel types underlies the diverse firing behavior of various types of neurons.

### Conclusions

We conclude that Y1eGFP neurons are excitatory interneurons that mainly exhibit phasic firing rather than delayed firing patterns, though delayed firing can be unmasked when current clamp recordings are initiated at hyperpolarized conditions that could be mimicked by Y1 receptor activation. Our results emphasize the importance of initial membrane potential when segregating SDH neurons by firing pattern.

In contrast to the vast majority of randomly sampled SDH neurons which do not exhibit spike frequency adaptation, a substantial subpopulation of Y1eGFP DSLF neurons were rapidly adapting. The incidence of rebound spiking was surprising high in Y1eGFP DSLF neurons, suggestive of their enrichment with T-type calcium channels that contribute to membrane hyperexcitability and nociception. We also found that actions potentials in Y1eGFP neurons, particularly of the DSLF type, could be elicited by electrical stimulation at a frequency and intensity sufficient to recruit C- and/or Aδ fibers, both of which are essential for the transmission of noxious somatosensation (Todd & Koerber, 2006; Yasaka *et al*., 2010). Further studies are needed to better characterize these inputs, as has been done for GRP^cre^-expressing neurons in the dorsal horn (Sun *et al*., 2017). Indeed, as discussed and illustrated in our recent review, we hypothesize that Aβ fibers provide inputs to Y1-expressing interneurons which may be particularly important to neuropathic pain in the setting of peripheral nerve injury (Nelson & Taylor, 2020). It remains to be determined whether NPY engages such mechanisms to exert its well-described actions to reduce behavioral signs of neuropathic pain (Smith *et al*., 2007; Nelson & Taylor, 2020).

## Supporting information

Statistical summary table

## Acknowledgements

We are grateful to Michael Gold for insightful discussions during the preparation of the manuscript. This study was supported by NIH Grants R01DA37621, R01NS45954, and R01NS62306 to BKT.

## Data availability statement

The data that support the findings of this study are available from the corresponding author upon reasonable request.

## Supporting information

The following additional supporting information may be found online in **“*Supporting Information for Online Publication*”:**

1. Statistical Summary Document is an excel spreadsheet listing all statistical outputs for Tables 2-5, 8 and Figures 10M-N and 11.
2. Tables of statistical results with data averaged across numbers of cells (small n) instead of numbers of mice (large N): Tables S2 – S5 correspond to Tables 2 – 5, respectively).
3. Figure 3S (Figure 3, Panel D), Figure 4S (Figure 4, Panel D), Figure 10S (Figure 10, Panels M-N), Figure 11S (Figure 11, Insets of A-B and panels E-F) and Figure 12S (Figure 12, Panel E) describes data with statistical analysis conducted on number of neurons (small n) rather than number of animals (large N).

