## Supplemental Tables and Figures for "Fast A-type currents shape a rapidly adapting form of delayed short latency firing of excitatory superficial dorsal horn neurons that express the NPY Y1 receptor"

**Supplementary Tables and Figures**

**Supplementary Tables**:

|  | Random | Y1eGFP |
| --- | --- | --- |
| Membrane capacitance (pF) | 28 ± 9 (32) | 29 ± 9 (76) |
| Input resistance (MΩ) | 703 ± 550 (32) | 707 ± 366 (76) |
| RMP, mV | -64 ± 7 (28) | -62 ± 9 (65) |
| AP threshold (mV) | -38.3 ± 5.6 (29) | -32.7 ± 5.1 (51)*** |
| AP width at base (ms) | 1.9 ± 0.5 (35) | 2.3 ± 0.4 (54)** |
| AP height from base (mV) | 82 ± 11 (35) | 83 ± 10 (54) |
| AHP amplitude (mV) | 20.4 ± 6.9 (35) | 20.4 ± 4.5 (54) |

**Table S2:** Passive membrane properties [membrane capacitance, input resistance, and resting membrane potential (RMP)] and active membrane properties [action potential (AP) threshold, width, height, and afterhyperpolarization (AHP)] of randomly sampled and Y1eGFP SDH neurons. Data were collected from 14 WT mice (Random) and 27 Y1eGFP mice. Values represent mean ± SD. Numbers of neurons are indicated in parentheses. **p<0.01 or ***p<0.001, Student’s t-test, between respective injecting strengths, Random vs. Y1eGFP.

|  | | Random | | |  | | Y1eGFP | | |
| --- | --- | --- | --- | --- | --- | --- | --- | --- | --- |
|  |  | LCI^A^ | MCI^B^ | HCI^C^ |  | LCI^D^ | | MCI^E^ | HCI^F^ |
| Spike latency (ms) | DSLF-N | 124 ± 41 (28) | - | - | DSLF-A | 103 ± 31 (26) *(AD) | | - | - |
|  | DLLF | 222 ± 36 (5) | - | - | DLLF | 197 ± 36 (10) | | - | - |
| No. of spikes | DSLF-N | 7.0±4.6 (25) | 19.8±14.0 (24) | 18.6±13.7 (25) | DSLF-A | 4.4±1.7 (33) **(AD) | | 8.4±5.8 (29) ***(BE) | 6.8±4.3 (32) ***(CF) |
|  | DLLF | 8±6 (8) | 15±10 (8) | 12±11 (8) | DLLF | 6±3 (23) | | 13±7 (23) | 11±6 (23) |
|  | IBF | 14±10 (5) | 30±29 (5) | 28±25 (5) | IBF | 3±1 (12) **(AD) | | 7±6 (11) *(BE) | 6±3 (12) *(CF) |
|  | TF | 5±2 (2) | 61±10 (2) | 70±3 (2) | TF | 8±1 (2) | | 74±47 (2) | 61±6 (2) |

**Table S3**: First spike latency in delayed firing neurons and number of spikes at different current injecting strengths from conditioning hyperpolarized potentials for randomly sampled (Random) and Y1eGFP SDH neurons. The latency to first spike was determined by injecting currents just above threshold (LCI). Numbers in brackets indicates number of neurons. LCI, MCI and HCI denote Low, Medium and High strengths of current injection. DSLF stand for delayed short latency firing (N is non-adapting, A is adapting), DLLF for delayed long latency firing, IBF for initial burst firing and TF for tonic firing. Values represent mean ± SD. *p<0.05, **p<0.01 or ***p<0.001, Student’s t-test, for respective injecting strengths, Random vs. Y1eGFP.

|  | | Random | | |  | Y1eGFP | | |
| --- | --- | --- | --- | --- | --- | --- | --- | --- |
|  |  | LCI^A^ | MCI^B^ | HCI^C^ |  | LCI^D^ | MCI^E^ | HCI^F^ |
| Initial spike frequency (Hz) | DSLF-N | 14±7 (23) | 34±14 (22) | 41±17 (23) | DSLF-A | 18±8 (34) | 30±9 (29) | 36±10 (32) |
|  | DLLF | 30±27 (7) | 49±32 (7) | 56±30 (7) | DLLF | 15±6 (22) *(AD) | 28±15 (22)  *(BE) | 32±15 (22) **(CF) |
|  | IBF | 31±21 (5) | 55±31 (5) | 74±34 (5) | IBF | 76±34 (12) *(AD) | 102±28 (11) **(BE) | 96±34 (12) |
|  | TF | 10±1 (2) | 74±18 (2) | 104±24 (2) | TF | 25±24 (2) | 88±76 (2) | 114±77 (2) |
| Average spike frequency (Hz) | DSLF-N | 14±8 (25) | 35±14 (24) | 42±18 (25) | DSLF-A | 15±7 (32) | 25±7 (29)  **(BE) | 30±7 (30) **(CF) |
|  | DLLF | 21±18 (8) | 32±13 (8) | 38±19 (7) | DLLF | 13±3 (21) *(AD) | 25±10 (21) | 29±11 (21) |
|  | IBF | 23±13 (5) | 46±22 (5) | 62±20 (5) | IBF | 54±26 (12) *(AD) | 62±23 (11) | 64±28 (11) |
|  | TF | 9±1 (2) | 61±10 (2) | 82±13 (2) | TF | 28±27 (2) | 75±47 (2) | 88±45 (2) |

**Table S4:** Characteristic firing frequencies (initial and average) at different current injection strengths from conditioning hyperpolarized potentials for randomly sampled (Random) and Y1eGFP SDH neurons. Numbers in brackets indicate number of neurons. LCI, MCI and HCI denote Low, Medium, and High strengths of current injection. DSLF stand for delayed short latency firing (N is non-adapting, A is adapting), DLLF for delayed long latency firing, IBF for initial burst firing and TF for tonic firing. Values represent mean ± SD. *p<0.05, **p<0.01 or ***p<0.001, Student’s t-test, for respective injecting strengths, Random vs. Y1eGFP.

|  | | Random | | |  | Y1eGFP | | |
| --- | --- | --- | --- | --- | --- | --- | --- | --- |
|  |  | LCI^A^ | MCI^B^ | HCI^C^ |  | LCI^D^ | MCI^E^ | HCI^F^ |
| Initial frequency adaptation | DSLF-N | -0.14±0.29 (25) | 0.03±0.39 (24) | 0.01±0.09 (25) | DSLF-A | -0.24±0.14 (34) | -0.12±0.11 (30) *(BE) | -0.12±0.09 (33) *(CF) |
|  | DLLF | 0.00±0.71 (8) | 0.09±0.29 (8) | -0.09±0.13 (8) | DLLF | -0.13±0.22 (23) | 0.13±0.24 (23) | 0.10±0.24 (23) *(CF) |
|  | IBF | 0.03±0.33 (5) | -0.16±0.24 (5) | -0.17±0.28 (5) | IBF | -0.46±0.10 (12) ***(AD) | -0.29±0.11 (11) | -0.22±0.29 (12) |
|  | TF | -0.09±0.01 (2) | 0.01±0.01 (2) | -0.04±0.02 (2) | TF | 0.33±0.12 (2) *(AD) | 0.04±0.13 (2) | -0.01±0.05 (2) |
| average frequency adaptation | DSLF-N | -0.02±0.23 (22) | -0.00±0.09 (23) | -0.03±0.07 (24) | DSLF-A | -0.11±0.04 (22)  *(AD) | -0.08±0.05 (26) ***(BE) | -0.08±0.05 (28) **(CF) |
|  | DLLF | -0.04±0.10 (8) | -0.03±0.04 (8) | -0.05±0.04 (7) | DLLF | -0.09±0.13 (20) | -0.03±0.04 (22) | -0.04±0.07 (22) |
|  | IBF | -0.01±0.12 (5) | -0.01±0.06 (5) | -0.03±0.05 (5) | IBF | -0.2±0.16 (7) *(AD) | -0.19±0.12 (9) *(BE) | -0.12±0.08 (11) *(CF) |
|  | TF | -0.42±0.43 (2) | -0.00±0.00 (2) | -0.00±0.00 (2) | TF | -0.15±0.17 (2) | -0.00±0.00 (2) | -0.01±0.01 (2) |
|  | TF | -0.42±0.43 (2) | -0.00±0.00 (2) | -0.00±0.00 (2) | TF | -0.15±0.17 (2) | -0.00±0.00 (2) | -0.01±0.01 (2) |

**Table S5**: Characteristic frequency adaptation [initial $\{{(f_{2}- f_{1})}/{f_{1}\}}$ and average of all $\{{(f_{i-1}- f_{i})}/{f_{1}\}}$] at different current injection strengths from conditioning hyperpolarized potentials for randomly sampled (Random) and Y1eGFP SDH neurons. Number in brackets indicates number of neurons. LCI, MCI and HCI denote Low, Medium and High strengths of current Injection. DSLF stand for delayed short latency firing (N is non-adapting, A is adapting), DLLF for delayed long latency firing, IBF for initial burst firing and TF for tonic firing. Values represent mean ± SD. *****p<0.05**,** **p<0.01 or ***p<0.001: Student’s t-test, for respective injecting strengths, randomly sampled vs. Y1eGFP.

**Supplementary Figures**:

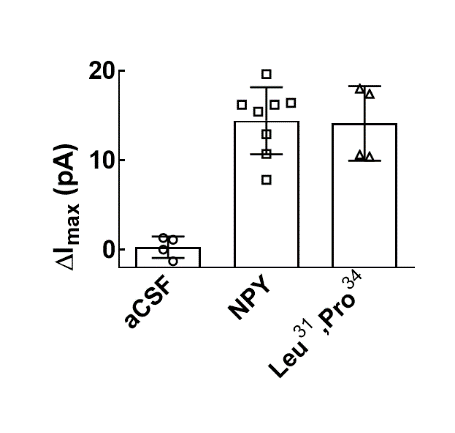

**Figure 3S.** Peak current amplitude within 10 s of application of aCSF (4 neurons from 3 mice), NPY (8 neurons from 4 animals responded out of a total of 34 Y1eGFP neurons given drug), or [Leu^31^,Pro^34^]-NPY (4 neurons from 3 animals responded out of a total of 11 neurons given drug). Values represent mean ± SD; each dot represents a neuron.

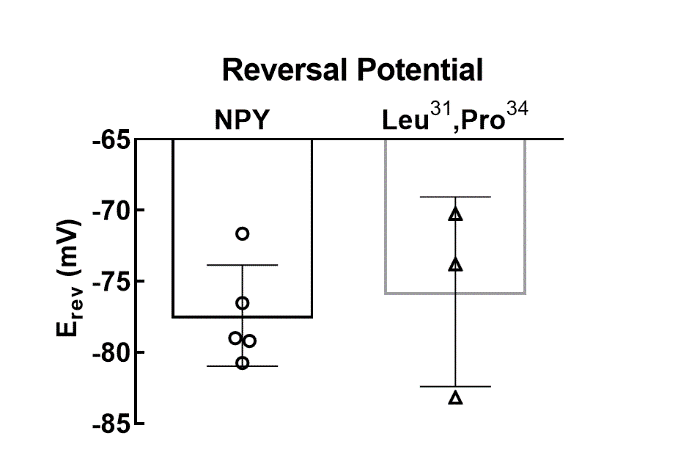

**Figure 4****S.** Mean reversal potentials [*E_rev_*] for current induced by 20 µM NPY (n=5 neurons, 3 animals) or 20 µM [Leu^31^,Pro^34^]-NPY (n=3 neurons, 3 animals). Values represent mean ± SD; each dot represents a neuron.

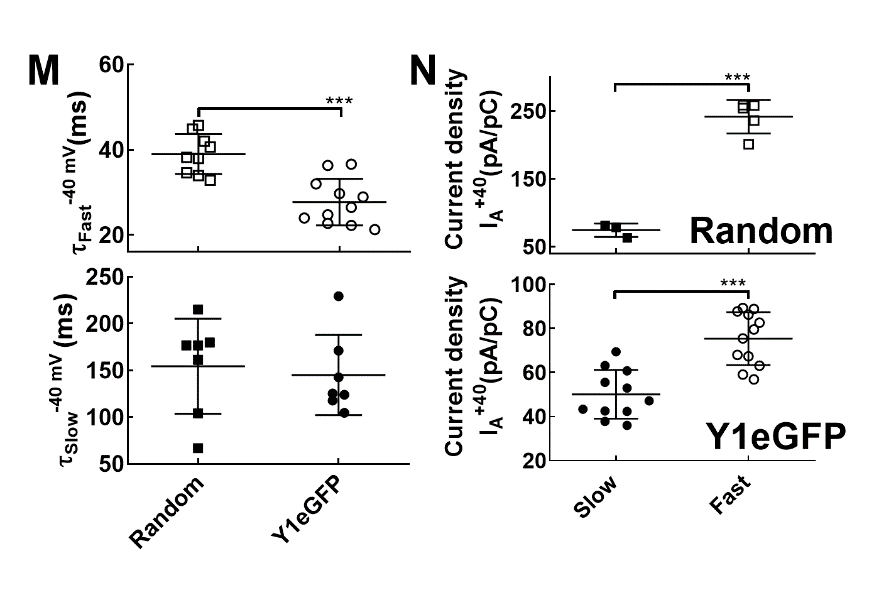

**Figure 10S.** (**M**) Decay constant obtained from Fast (top, RS: 8 neurons, 6 animals, Y1eGFP: 11 neurons, 8 animals; *** P=0.0003: Student’s t-test, Random vs Y1eGFP) and Slow (bottom; RS and Y1eGFP: 7 neurons, 6 animals) A-type currents in randomly sampled and Y1eGFP neurons in response to steps from -100 to -40 mV. *** P<0.001: Student’s t-test, Random vs Y1eGFP. (**N**) Current density calculated for Fast (RS: 5 neurons, 4 animals, Y1eGFP: 12 neurons, 11 animals) and Slow (RS: 3 neurons, 3 animals, Y1eGFP: 12 neurons, 11 animals) A-type currents in randomly sampled (top) and Y1eGFP (bottom) neurons. *** P<0.001: Student’s t-test, Slow vs. Fast. Values represent mean ± SD; each dot represents a neuron.

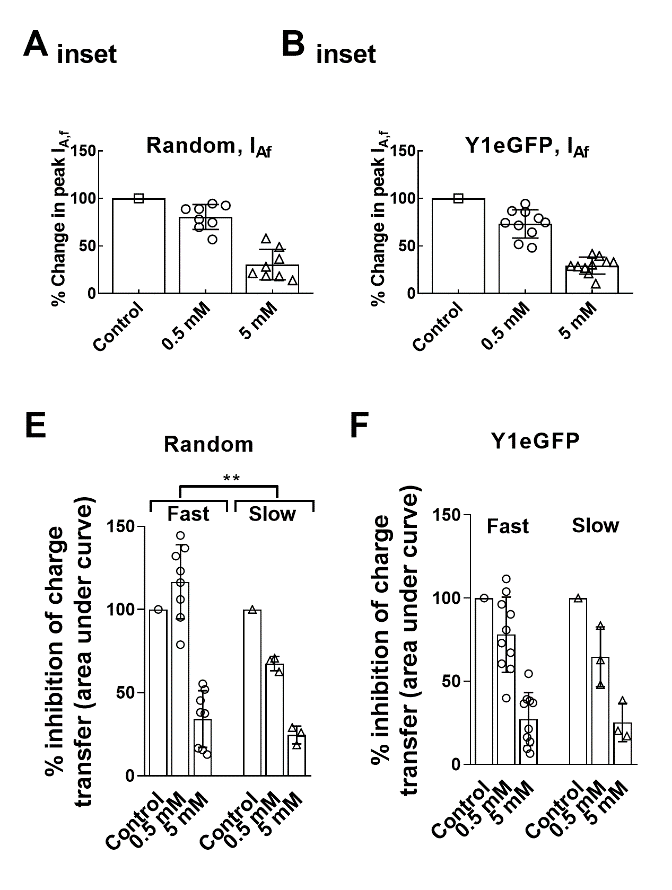

**Figure 11S. Insets of A-B**: Inhibition of the Fast and Slow A-type currents by 4-AP in SDH neurons. Percentage change in peak currents induced by 4-AP (RS: 8 neurons from 5 mice; Y1eGFP: 10 neurons from 4 mice). (**E-F**) Summary of inhibition by 0.5 mM and 5 mM 4-AP to Fast (I_Af_) and Slow (I_As_) A-type currents, expressed as percentage change in area under the curve in randomly sampled (**E**) and Y1eGFP (**F**) neurons. Values represent mean ± SD; each dot represents a neuron. **p=0.005: 2-way ANOVA, 4-AP conc. vs A-current type.

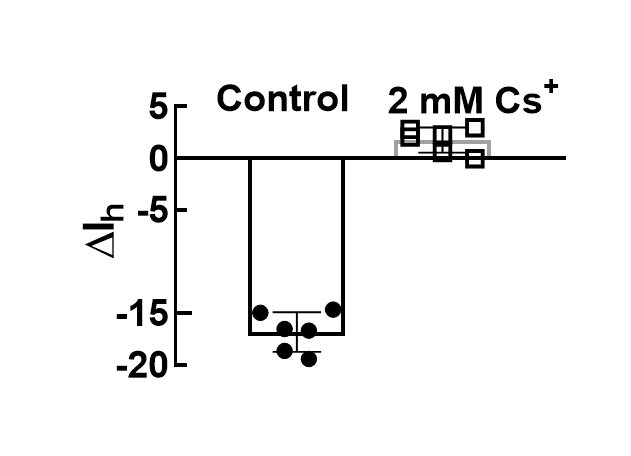

**Figure 12S.** Quantification of the relative (completely) block by 2 mM Cs^+^ of *I_h_* in randomly sampled neurons (n=6 neurons, 5 animals). Values represent mean ± SD; each dot represents a neuron.
